# Prediction of protein-RNA interactions from single-cell transcriptomic data

**DOI:** 10.1101/2023.07.12.548714

**Authors:** Jonathan Fiorentino, Alexandros Armaos, Alessio Colantoni, Gian Gaetano Tartaglia

## Abstract

RNA-binding proteins play a crucial role in regulating RNA processing, yet our understanding of their interactions with coding and non-coding RNAs remains limited. Experimental studies are typically restricted to a small number of cell lines and a limited set of RNA-binding proteins (RBPs). Although computational methods based on sequence and structure can predict protein-RNA interactions accurately, they often lack the ability to consider cell-type-specific gene expression and the broader context of gene regulatory networks (GRNs).

In the present study, we assess the performance of several GRN inference algorithms in predicting protein-RNA interactions from single-cell transcriptomic data, and we propose a pipeline, called scRAPID (single-cell transcriptomic-based RnA Protein Interaction Detection), that integrates these methods with the *cat*RAPID algorithm, which can identify direct physical interactions between RBPs and RNA molecules.

Our approach demonstrates that RBP-RNA interactions can be inferred from single-cell transcriptomic data, with performances comparable or superior to those achieved for the well-established task of inferring transcription factor-target interactions. Notably, the incorporation of *cat*RAPID significantly enhances the accuracy of identifying interactions, particularly with long non-coding RNAs, and enables the identification of hub RBPs and hub RNAs. Additionally, we show that interactions between RBPs can be detected based on their inferred RNA targets.

We have made the software freely available at https://github.com/tartaglialabIIT/scRAPID.

## Introduction

RNA-binding proteins (RBPs) are key players in post-transcriptional regulation of gene expression (1), being involved in several aspects of RNA processing, including polyadenylation, splicing, capping and cleavage. They bind both coding and non-coding RNAs through RNA-binding domains, although several unconventional modes through which RBPs recognize their targets have been recently characterized (2). Recent advances in high-throughput experimental techniques, such as enhanced Crosslinking Immunoprecipitation (eCLIP) (3), provided a large catalog of known interactions of RNA with RBPs (4). However, current knowledge based on CLIP-Seq data is limited due to two main reasons: i) the experiments were performed in few cell lines, but emerging evidence indicates that RBP-RNA interactions occur specifically in distinct cell types and at determined time points (5, 6); ii) despite CLIP-Seq data are available for hundreds of RBPs, thousands of them are known at present and their list is continuously expanded in multiple species (2, 7). Moreover, the detection efficiency of CLIP-based techniques has several limitations, both in terms of sensitivity (e.g., low-abundance transcripts could be difficult to detect due to low crosslinking efficiency), specificity (8, 9) and reproducibility (10).

In parallel, several computational approaches, trained on the available experimental data, have been developed for the prediction of protein-RNA interactions (11–13). We previously developed the *cat*RAPID method, which combines information from the secondary structure, hydrogen bonding and van der Waals contributions to estimate the interaction propensity of protein-RNA pairs with an accuracy of 78% or higher (14, 15). The interaction propensity calculated with *cat*RAPID correlates with the experimental binding affinities (11, 16, 17) and was successfully exploited to identify the binding partners of non-coding transcripts such as *Xist (15)*, *HOTAIR (18)*, HOTAIRM1 (19) and *SAMMSON* (20), as well as the interactomes of RNA genomes (21).

The identification of common targets of RBPs holds significant importance, especially considering the involvement of specific RNAs such as Xist (22) and Neat1 (23) in promoting the formation of liquid-liquid phase-separated organelles. RBPs have been found to assemble in such organelles, including stress granules, which play a crucial role in regulating gene expression (24–26). In some cases, aberrant macromolecular assembly and dysregulation of these organelles have been associated with neurodegenerative disease (27). However, the impact of these changes in interaction networks on aberrant macromolecular assembly is still an unexplored area of research.

In the last decade, single-cell RNA-sequencing (scRNA-seq) provided an unprecedented resolution of the cell type composition and transcriptional landscape of many organs and tissues in different organisms, leading to global atlas projects such as the Human Cell Atlas (28, 29) and the Mouse Cell Atlas (30). Beyond the identification of new cell types and subtypes, statistical and machine-learning methods for the inference of Gene Regulatory Networks (GRNs) from scRNA-seq data have been developed, with the goal of identifying the complex regulatory interactions between transcription factors (TFs) and their targets (31, 32).

Despite the centrality of RBPs in the cellular regulatory machinery, transcriptome-wide experimental identification of RBP targets at single-cell resolution is still in its infancy (6). Notably, the interaction propensity of an RBP with its target RNA is associated with a correlation between their expression levels, as shown by computational analyses of RBP-RNA interactions and bulk RNA-seq data (33), and RNA and protein expression levels are highly correlated for RBPs (34, 35). Furthermore, recent work has shown that it is possible to identify RBP regulators through the integration of scRNA-seq data and RNA-binding motif information (36).

In this study, we conducted a systematic evaluation of GRN inference methods for accurately predicting protein-RNA interactions using single-cell transcriptomic data. We propose a pipeline, called scRAPID, which integrates these methods with *cat*RAPID to enhance the inference performance. To assess their behavior in comparison to the classical task of TF-target inference, we focused on two cell lines (HepG2 and K562) with publicly available Chromatin Immunoprecipitation and sequencing (ChIP-seq), enhanced Crosslinking and Immunoprecipitation (eCLIP), bulk RNA-seq, and scRNA-seq data.

Initially, we demonstrated that scRNA-seq data can be effectively used to infer protein-RNA interactions, exhibiting performances comparable to or even surpassing those achieved for TF-target inference. Subsequently, we improved the performance by employing *cat*RAPID predictions (14, 15) to filter the returned GRNs from each method. Leveraging RNA-seq data obtained from experiments involving the knockdown of RNA-binding proteins (RBPs), specifically pooled short-hairpin RNA sequencing (shRNA RNA-seq), we demonstrated the efficacy of *cat*RAPID in filtering out indirect interactions.

Furthermore, we assessed the performance of the methods in predicting interactions between RBPs and long non-coding RNAs (lncRNAs). Despite the limited availability of experimental data for RBP-lncRNA interactions compared to mRNA interactions, we consistently achieved superior performance in inferring RBP-lncRNA interactions, particularly with scRNA-seq datasets obtained through the latest full-length sequencing protocols such as STORM-seq (37) and Smart-seq3 (38).

Additionally, we evaluated the ability of the inference methods to identify hub RBPs, hub mRNAs, and hub lncRNAs, which are defined as regulators of a large number of RNAs or as RNAs regulated by a large number of RBPs, respectively. Notably, recent benchmarking studies have demonstrated that even when GRN inference methods achieve moderate performance in predicting the edges of the ground truth network, they often excel in predicting hub genes, which typically serve as master regulators in the biological processes under investigation (39). Confirming these findings within the context of protein-RNA interactions, we also observed that *cat*RAPID enhanced the performance of the inference methods in identifying network hubs.

To validate our methodology across different organisms and experimental techniques beyond eCLIP, we analysed a mouse cell line that recapitulates myoblasts-to-myotubes differentiation and retinoic acid-driven differentiation of mouse embryonic stem cells. Specifically, we evaluated the inference performance for two RBPs, ADAR1 and Caprin1, which play a role in stress granule formation (40, 41) and for which RNA targets are available from RNA immunoprecipitation followed by RNA-sequencing (RIP-seq) experiments. Our results showcased favorable performance across most of the methods, with *cat*RAPID consistently improving the predictive accuracy.

Lastly, we demonstrated the feasibility of predicting direct RBP-RBP interactions by leveraging the overlap of RNA targets inferred from the scRNA-seq data.

In summary, our study presents a novel and scientifically elegant evaluation of GRN inference methods for predicting protein-RNA interactions from single-cell transcriptomic data. By introducing *cat*RAPID and conducting extensive validations, we significantly enhance the inference performance and provide valuable insights into RBP-lncRNA interactions, hub identification, and direct RBP-RBP interactions.

## Methods

We used human hepatocellular carcinoma (HepG2) and human lymphoblastoma (K562) cell lines in most of the analyses since, for these cell types, ChIP-seq, eCLIP and shRNA RNA-Seq datasets are available from the ENCODE project (42, 43), and multiple scRNA-seq datasets obtained through different protocols are available from public repositories.

### Single-cell RNA-seq datasets

#### HepG2

For the HepG2 cell line we selected three scRNA-seq datasets, obtained through the Smart-seq2, DNBelab C series Single-cell System, a droplet-based system similar to that from 10x Genomics in cell throughput and data formatting, and SCAN-seq2 (a single-cell Nanopore-based sequencing protocol) sequencing protocols.

The Smart-seq2 dataset is available on the Gene Expression Omnibus (GEO) (44) under accession number GSE150993 (45). We selected only the 68 live cells in the dataset.

The DNBelab dataset is available on GEO under accession number GSM5677000 (46). It contains 1628 cells.

The SCAN-seq2 dataset is available on GEO under accession number GSE20356 (47). We selected the “9CL” library, which contains 80 HepG2 cells.

#### K562

For the K562 cell line we selected five scRNA-seq datasets, obtained through the CEL-seq, STORM-seq, Smart-seq3 and SCAN-seq2 sequencing protocols.

The CEL-seq dataset is available on GEO under accession number GSM1599500 (48). It contains 239 cells.

The STORM-seq dataset is available on GEO under accession number GSE181544 (37). It contains 70 cells. The authors provided 3 processed datasets obtained from the same cells with different sequencing depth (100k, 500k and 1M reads). We used the dataset with highest depth (1M) in all the analyses.

The Smart-seq3 dataset is available on Arrayexpress under accession number E-MTAB-11467 (38, 49). It contains 231 cells sequenced with different reaction volumes (1 μl, 2 μl, 5 μl and 10 μl) and with cDNA clean-up or dilution. We performed a standard pre-processing using the R package Seurat v4.1.0 (50), then we used the function “FindMarkers” to find differentially expressed genes between the reaction volumes of 1 μl and 10 μl. We found only 1 gene with adjusted p-value smaller than 0.05. Instead, we found 32 differentially expressed genes between the “cleanup” and “diluted” condition. We kept all 231 cells for downstream analysis but we removed the 32 identified differentially expressed genes.

The SCAN-seq2 datasets for the K562 cell line are provided in the same study mentioned above for HepG2. We selected two libraries in this case: the “9CL”, containing 159 cells, and the “UMI200”, since it is the library with the highest sequencing depth, containing 96 cells.

#### Pre-processing and gene selection

We used Scanpy (version 1.8.2) (51) for the pre-processing and gene selection steps of the scRNA-seq datasets analysis. We removed spike-in genes where present. We filtered out genes expressed in less than 10% of the cells (1% for the HepG2 DNBelab dataset that has 1628 cells) using the function “scanpy.pp.filter_genes’’ and we removed mitochondrial genes. We used the TPM matrices for the HepG2 Smart-seq2 and the K562 STORM-seq datasets; we log-transformed them using the function “scanpy.pp.log1p”. We normalized the UMI counts for the HepG2 DNBelab, K562 CEL-seq, K562 Smart-seq3 and SCAN-seq2 datasets using the function “scanpy.pp.normalize_total” and we log-transformed the normalized counts using the function “scanpy.pp.log1p”.

Two inference algorithms (SINCERITIES and TENET) require cells ordered in pseudotime. To this end, we computed the top 2000 highly variable genes using the function “scanpy.pp.highly_variable_genes” and we scaled the data to zero mean and unit variance using the function “scanpy.pp.scale”, clipping values larger than 10 (parameter “max_value=10”). We performed a principal component analysis using the function “scanpy.tl.pca”, with “svd_solver=arpack”. We computed a k-nearest neighbor graph (“scanpy.pp.neighbors”) and a UMAP (52) (“scanpy.tl.umap”). Next, we computed a diffusion map (53) (“scanpy.tl.diffmap”) and the diffusion pseudotime (54) (“scanpy.tl.dpt”) choosing the root cell based on the UMAP coordinates, since in these cell lines there is not an obvious starting cell for the pseudotime computation.

For the gene selection step, we followed the BEELINE evaluation framework (31) and selected the top 500 and 1000 highly variable genes in each dataset. We restricted the gene sets for selection to protein-coding and long non-coding RNAs, according to the annotation in Gencode V41 (55). We added to each dataset the highly variable transcription factors or the RNA binding proteins present in the eCLIP experiments from the ENCODE project. We used a list of 1563 transcription factors provided in (31).

Regarding the analyses involving comparisons between GRN inference on mRNAs and lncRNAs (**Figure 3** and **Supplementary Figures** reported in the paragraph ‘Predicting protein interactions with long non-coding RNAs’), we selected the top 400 highly variable mRNAs or lncRNAs in each dataset. We added to each dataset the eCLIP RBPs as shown before.

### Ground truth networks

#### TF-target

##### ChIP-seq

We downloaded ChIP-seq data for the HepG2 (56) and K562 cell lines from the ENCODE project portal (https://www.encodeproject.org/) (42). Accession codes and metadata obtained from the ENCODE project portal are reported in the **Supplementary Tables** linked to section ‘Protein-RNA interactions can be inferred from single-cell RNA-seq data’. Considering data associated with the GRCh38 assembly and experiments with multiple biological replicates, we selected 589 and 477 BED files with peaks merged using the IDR approach for HepG2 and K562, respectively. Next, using the “window” module from BEDTools (version 2.30.0) (57), we identified the target gene for each peak, defined as the closest gene whose transcription start site (TSS) is less than 50 kilobases away from the peak. TSS information was retrieved from the “upstream1000.fa” file provided by the UCSC Genome Browser (58) (available at https://hgdownload.soe.ucsc.edu/goldenPath/hg38/bigZips/), which reports the transcription starts of RefSeq genes (59) with annotated 5’ UTRs.

#### RBP-target

##### eCLIP

The BED files relative to the peaks identified in each biological replicate of eCLIP experiments conducted for 103 RBPs in HepG2 cells and 120 RBPs in K562 cells (3) were downloaded from the ENCODE project portal, selecting the GRCh38 assembly and “BED narrowpeak” file type. Accession codes and metadata obtained from the ENCODE project portal are reported in the **Supplementary Tables** mentioned in paragraph ‘Protein-RNA interactions can be inferred from single-cell RNA-seq data’. Next, we filtered the files of the single replicates with logFC > 1 and -log10 p-value > 3 and we took the intersection between the BED files of the single replicates for each RBP, using the “intersect” module from BEDTools (version 2.30.0). The same tool was used to find overlaps between the consensus peaks and the canonical isoforms of mRNA and lncRNA obtained from Ensembl 107 (60) distinguishing between exonic and intronic peaks. For the analyses in the manuscript we employed only the interactions involving exonic regions, because the inference of the interaction propensity of unspliced RNA molecules with *cat*RAPID is a computationally demanding task that would require an *ad hoc* fragmentation procedure to deal with very long sequences.

##### shRNA RNA-Seq

We used the metadata from (4) (file “’41586_2020_2077_MOESM4_ESM.xlsx’”, sheet name: “KD-RNA-seq”, column “RBP knockdown DESeq after batch Correction”) to obtain the tsv file names for each RBP. We downloaded the tsv files from the ENCODE project, and we retained only targets with FDR < 0.05.

To identify indirect RBP-RNA interactions (**Figure 2C** and **Supplementary Figures** reported in paragraph ‘*cat*RAPID improves prediction performance and removes indirect interactions’), for each cell line we considered the interactions involving RBPs present in both eCLIP and shRNA RNA-Seq datasets (92 RBPs for HepG2 and 110 for K562), and, for each RBP, we removed the eCLIP-identified targets from those detected *via* shRNA RNA-Seq. After this filter we obtained 259327 interactions involving 92 RBPs in HepG2 and 176843 interactions involving 110 RBPs in K562.

### GRN inference methods and implementation

For the inference of GRNs from scRNA-seq data, we chose the three top performing methods from BEELINE (31) (PIDC, GRNBOOST2, SINCERITIES). We also added two more recent methods that have been shown to outperform previous ones (TENET and DeePSEM), and a method not specifically designed for scRNA-seq, but that has good performance and it is widely used (ARACNe).

#### PIDC

PIDC is a GRN inference algorithm based on multivariate information measures (61). Specifically, it uses Partial Information Decomposition (PID) between triplets of genes to find putative functional interactions between genes. It outputs an undirected network.

#### GRNBOOST2

GRNBoost2 uses stochastic gradient boosting regression to select the top regulators for each gene in the dataset (62). It is based on GENIE3 (63), a regression method initially designed for bulk transcriptomic data, but it is faster, thus it is more suited for scRNA-seq data. It outputs a directed network.

#### SINCERITIES

SINCERITIES requires cells ordered in pseudotime. It computes temporal changes in the expression of each gene in pseudotime using the Kolmogorov-Smirnov statistic (64). It uses Granger causality to infer connections between regulator and target genes. The GRN inference is formulated as a ridge regression problem. It outputs a signed and directed network; however, in this work we do not take into account the sign information.

#### TENET

TENET computes the transfer entropy, a measure of directed information transfer, between the expression profiles along pseudotime of each pair of genes in the dataset (65). Potential indirect interactions are trimmed applying the Data Processing Inequality (DPI). The False Discovery Rate (FDR) of the interactions is computed by performing a one-sided z-test considering the trimmed values of transfer entropy as normally distributed. TENET outputs a directed network. Due to the indirect interaction trimming, TENET outputs smaller networks compared to the other inference methods, making it more suited to be used on datasets with a larger number of genes. For this reason, we include TENET networks obtained in three different ways:

- TENET: Full network without indirect interaction trimming.
- TENET_A: Network obtained after indirect interaction trimming (cutoff = -0.1) and FDR < 0.01. This is the original usage.
- TENET_B: Network obtained after indirect interaction trimming (cutoff = -0.1) and FDR < 0.5.

For TENET implementation, we follow the installation and usage instructions provided at https://github.com/neocaleb/TENET.

#### DeePSEM

DeePSEM is a deep generative model designed for scRNA-seq data that can simultaneously infer a GRN, embed and visualize scRNA-seq data and simulate them (66). DeePSEM jointly models the GRN and the transcriptome by generating a Structural Equation Model (SEM) through a beta Variational Auto-Encoder (beta-VAE). Following the original implementation, we use the ensemble strategy to obtain more stable predictions, namely we train DeePSEM on the same dataset with ten different random initializations. The final GRN is obtained by averaging the adjacency matrices derived from the ten trained models. Following the implementation provided in https://github.com/HantaoShu/DeepSEM(GRN_inference_tutorial.ipynb), we used DeePSEM in cell-type specific mode (task = celltype_GRN). DeePSEM outputs a directed network.

#### ARACNe

ARACNe (Algorithm for the reconstruction of accurate cellular networks) is one of the first GRN inference methods based on information theory, initially designed for microarray data (67). It infers putative direct regulatory relationships between regulator and target genes using mutual information (MI). In a first filtering step, interactions with low MI are filtered out based on a threshold computed under the null hypothesis of independence of two genes. Subsequently, the DPI is applied to filter out indirect interactions. ARACNe outputs a directed network. In this work, we use a faster re-implementation of ARACNe based on Adaptive Partitioning (ARACNe-AP) (68), since it is more suited to deal with the larger number of samples in scRNA-seq data. We follow the steps for its implementation provided in https://github.com/califano-lab/PISCES/ (69), in the folder /tree/master/data. We ran ARACNe-AP on a HPC cluster using a Singularity container that we built. For datasets with more than 1000 cells, we ran ARACNe-AP on 250 metacells computed following the instructions in https://github.com/califano-lab/PISCES/blob/master/vignettes/general-workflow.Rmd.

#### Implementation of the GRN inference

GRN inference is based on BEELINE (31), whose installation instructions and documentation are available at https://murali-group.github.io/Beeline/. PIDC, GRNBOOST2 and SINCERITIES were already available in BEELINE, thus we used the Docker containers provided in it. For TENET, we followed the instructions provided at https://github.com/neocaleb/TENET, as described before, and we used custom bash and Python scripts, which we provide in our Github repository (https://github.com/tartaglialabIIT/scRAPID), to include it in the BEELINE pipeline. DeePSEM instead runs on a GPU architecture and it was implemented following the instructions provided by the authors (https://github.com/HantaoShu/DeepSEM). ARACNe-AP was run on a HPC cluster following the instructions provided by the authors (https://github.com/califano-lab/PISCES/tree/master/data). We provide the “def” file for building a Singularity image for running ARACNe-AP and custom bash scripts in our Github repository. Finally, we used a custom Python script to format the results of all the GRN inference methods as in BEELINE.

#### *cat*RAPID

*cat*RAPID is an algorithm that computes an interaction propensity score between a protein and a RNA based on their sequence, using information from the secondary structure, hydrogen bonding and van der Waals contributions (14, 15). Canonical protein sequences in FASTA format for the RBPs used in this study were obtained from Uniprot (70). Regarding the RNAs, we used the sequence of the canonical isoforms retrieved from Ensembl (version 107).

For the computation of the interaction propensity scores, we followed the fragmentation-based approach of the “*cat*RAPID fragment” module (71), also used in *cat*RAPID omics v2.0 (17) and RNact (72). The final interaction propensity for a protein-RNA pair is defined as the maximum over the distribution of the interaction propensities of the fragments, as in RNAct (72).

To facilitate the usage of scRAPID with new scRNA-seq datasets in different organisms, we provide a SQL database containing the maximum interaction propensity scores from *cat*RAPID for: i) 3131 RBPs vs 62055 RNAs (all the canonical isoforms for the full transcriptome) in human, for a total of 194.3 millions interactions; ii) 2900 RBPs and 53087 RNAs (all the canonical isoforms for the full transcriptome) in mouse, for a total of 154.0 millions interactions. The lists of human and mouse RBPs were compiled by combining the RBPs from the RBP2GO database having score larger than 10 (7) with those that make up the *cat*RAPID *omics* v2.0 RBP libraries (17); the latter sets were further expanded by including, for human and mouse, proteins that are orthologous to the RBPs identified in mouse and human, respectively.

The SQL database can be queried via “curl”; further details and example queries are provided in our Github repository (https://github.com/tartaglialabIIT/scRAPID).

Interactions that are missing in our database, for instance involving organisms other than human and mouse or RNA isoforms other than the canonical one in human and mouse, can be computed using the *cat*RAPID *omics* v2.0 web server (http://service.tartaglialab.com/page/catrapid_omics2_group).

### RBP co-interaction analysis

#### scRNA-seq datasets

##### HEK293T

We selected two scRNA-seq datasets for the HEK293T cell line. The 10x dataset is available from the website of 10x Genomics (73) and it contains 2885 cells. The Smart-seq3 dataset is available on ArrayExpress with accession E-MTAB-8735 (38) and the single cell identifier column in the sample information table is “HEK293T Smart-seq3”; it contains 117 cells. We followed the same pre-processing steps explained above for the HepG2 and K562 datasets.

##### HCT116

The scRNA-seq dataset for the HCT116 cell line is available on GEO with accession number GSE149224 (74). The dataset includes three different cell lines (RKO, HCT116 and SW480) treated with different doses of 5-fluorouracil treatment to study the DNA-damage response of the transcriptome.

We selected only the 3011 HCT116 cells and we exploited the presence of treated cells to compute the diffusion pseudotime. We followed the pre-processing steps explained above for the HepG2 and K562 datasets, then we computed a diffusion map of HCT116 cells. We computed the diffusion pseudotime choosing as the root cell the control cell farthest from the treated ones. Next, we kept only the 2161 control cells for downstream analysis, following the analyses done for the HepG2 and K562 cell lines for gene selection and GRN inference.

#### RBP co-interaction prediction

RBP co-interactions were predicted based on the overlap of the RNA targets inferred by each GRN inference algorithm. We considered as RBPs the intersection between human RBPs present in the RBP2GO database (7), with RBP2GO score larger than 10, and the proteins present in the BioPlex Interactome database (75), which contains protein-protein interactions measured with Affinity Purification Mass Spectrometry (AP-MS), for the corresponding cell line (HEK293T or HCT116). We obtained a list of 1808 and 1509 RBPs for the HEK293T and HCT116 cell lines, respectively, including 12730 and 9700 BioPlex interactions.

For HEK293T, 186, 363 and 562 RBPs were included in the top 1000, 2000, 3000 HVGs selected from the 10x scRNA-seq dataset, respectively; 49, 145 and 247 RBPs were included in the top 1000, 2000, 3000 HVGs selected from the Smart-seq3 scRNA-seq dataset, respectively.

For HCT116, 164, 341 and 520 RBPs were included in the top 1000, 2000, 3000 HVGs selected from the Drop-seq scRNA-seq dataset, respectively.

Next, for the GRN inference methods that output more than the 5% of the possible edges, we cut the ranking to this threshold. For the other methods we kept all the edges returned. Then, for each pair of RBP, we computed the Jaccard coefficient between their sets of targets.

We ranked the RBP-RBP pairs according to the value of the Jaccard coefficient and we ran a Gene Set Enrichment Analysis for each GRN inference algorithm and scRNA-seq dataset (with 1000, 2000 and 3000 HVGs selected) using the R package “fgsea” (76). The ground truth interactions are those obtained from the BioPlex Interactome database for each cell line. The code to predict and evaluate RBP co-interactions is provided in our Github repository https://github.com/tartaglialabIIT/scRAPID.

Additional information about **Methods** is in the **Supplementary Materials**.

## Results

### Protein-RNA interactions can be inferred from single-cell RNA-seq data

A recent work provided a framework, called BEELINE, to evaluate the performances of algorithms for the prediction of GRNs from scRNA-seq data (31). The main result of the study is that, when it comes to predicting ChIP-Seq derived cell type-specific TF-target interactions, the performance of 12 algorithms is generally moderate (31).

Thus, to assess the performances of such algorithms in predicting RBP-RNA interactions, and to compare them with those obtained for the TF-target inference task, we selected datasets for two cell lines, HepG2 and K562, for which both ChIP-seq and eCLIP data are available from the ENCODE project (42, 43). We used 8 publicly available scRNA-seq datasets obtained with different sequencing protocols, including both full-length and 3’ end-based protocols; their characteristics are provided in **Supplementary Table S1**, while further technical details about the datasets and their pre-processing are provided in the **Methods** section.

Next, we selected six GRN inference algorithms, including the top three performing ones from BEELINE (PIDC (61), GRNBOOST2 (62) and SINCERITIES (64)), two recent methods that were shown to outperform the methods used in BEELINE (TENET (65) and DeePSEM (66)) and ARACNe (67), a method initially designed for bulk RNA-seq, but that has been widely used on scRNA-seq data with appreciable performance (77). The methods use different statistical models and theories to infer regulatory interactions from scRNA-seq data; details about their features and implementation are provided in the **Methods** section. For TENET, we evaluated three different inferred GRN types: the full inferred network without any filtering (indicated as TENET in the figures) and the networks on which we applied the data processing inequality (DPI) for removing indirect interactions, followed by the application of a stringent (TENET_A) or loose (TENET_B) threshold on the False Discovery Rate (FDR) (see **Methods** for further details).

Following BEELINE, we selected the top 500 or 1000 highly variable genes (HVGs) for each dataset, then we added the highly variable TFs or the RBPs for which an eCLIP experiment is present in the ENCODE project for the corresponding cell line (**Supplementary Table S2**). We ran the inference methods on each dataset and we measured the inference performance on TF-target or RBP-target interactions using the Early Precision Ratio (EPR) (31), which is the fraction of true positives in the top *k* edges of the inferred network, where *k* is the number of edges in the ground truth network, divided by the density of the ground truth network (see **Supplementary Materials** for details). In **Figure 1A** we show the probability density of the EPR values for all the datasets and algorithms with 500 HVGs selected, for TF-target and RBP-target interactions. The performances for the TF-target inference task, in which the inferred networks are compared to cell-type specific ChIP-seq data, are in line with those observed by previous studies in other single-cell datasets and they are slightly better than a random prediction (black dashed line) for all datasets and methods (**Figure 1A**).

**Figure 1.**
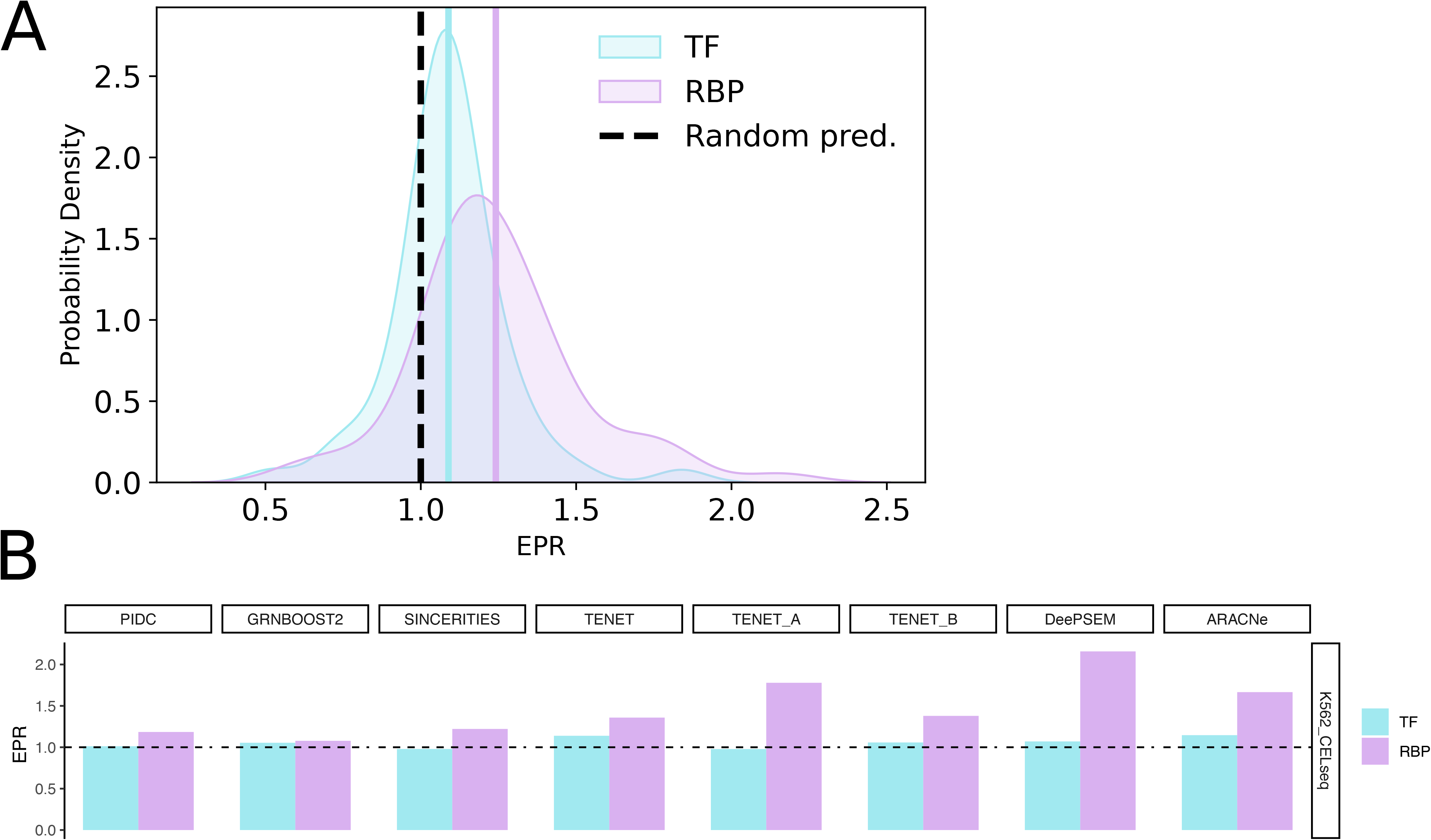
Performances obtained for the prediction of TF-target and RBP-target interactions from scRNA-seq data. (**A**) Probability densities of the Early Precision Ratio (EPR) measured across methods and datasets for the TF-target and RBP-target datasets. The ground truth network is given by cell-type specific ChIP-seq and eCLIP interactions for TF-target and RBP-target interactions, respectively (p-value= 4.4 x 10^-5^, Kolmogorov-Smirnov test). (**B**) Bar plots showing the EPR obtained for TF-target and RBP-target interactions by each GRN inference method, for the K562 CEL-seq scRNA-seq dataset. Both panels refer to the analyses performed using the top 500 HVGs. The black dashed line shows the EPR of a random predictor.

By contrast, the overall performance for RBP-target interactions, evaluated on cell-type specific eCLIP data, is significantly higher (p=4.4 x 10^-5^, Kolmogorov-Smirnov test, **Figure 1A**). The datasets with 1000 HVGs show similar results (**Supplementary Figures S1 and S2A**). **Figure 1B** shows the performances of individual algorithms for the K562 dataset sequenced with the CEL-seq protocol, for which, on average, we obtained the best performances for the RBP-target inference task.

The results for the other scRNA-seq datasets are shown in **Supplementary Figures S1 and S2B**, for 500 and 1000 HVGs, respectively. Interestingly, the EPR of the RBP-target inferred interactions is larger than the EPR of the TF-target ones in 73.4% of the cases. The statistics of the ChIP-seq and eCLIP ground truth networks for each dataset are reported in **Supplementary Figure S3**, together with the heatmaps of EPR values. On average, ARACNe is the top performing method for the TF-target inference task, while the dataset in which these interactions are best predicted is HepG2 DNBelab, as confirmed also for the datasets with 1000 HVGs (**Supplementary Figures S2 and S3**). For the inference of RBP-target interactions, DeePSEM is the top method in terms of average EPR over the datasets when considering 500 HVGs (**Supplementary Figures S1 and S3**), while it has a drop in performance for the datasets with 1000 HVGs, in which the best method is TENET_A (**Supplementary Figures S2 and S3**). The large increase in performance of the latter algorithm is likely due to the larger number of interactions inferred on datasets with 1000 HVGs. Indeed, TENET_A is very strict in considering significant interactions (see **Methods** for details), leading to the elimination of the majority of interactions for some datasets in the setting with RBPs and 500 HVGs. Regarding the datasets, in the case of 500 HVGs K562 CEL-seq is the best for inferring RBP-RNA interactions (**Figure 1B**), while with 1000 HVGs the top dataset becomes K562 STORM-seq **(Supplementary Figure S2**), in particular thanks to the performance of TENET_A, which was not present for this dataset with 500 HVGs due to the small number of interactions returned by this method.

Our systematic evaluation shows for the first time that RBP-RNA interactions can be inferred from scRNA-seq data, with performance (in terms of EPR) similar to or better than the ones obtained for the “classical” TF-target inference task. However, it should be noticed that the percentages of true positives reach a maximum of ∼35% at the top of the ranking of inferred interactions (**Supplementary Figures S4 and S5**). This is likely due to the confounding presence of direct protein-protein and RNA-RNA interactions, and of indirect interactions, which might be hard to disentangle from each other just looking at the interdependence of RNA expression levels. Another reason for the slightly low true positive rate is the incompleteness of the ground truth eCLIP network, especially due to detection limits.

### *cat*RAPID improves prediction performance and removes indirect interactions

We reasoned that an improvement in the performance of the GRN inference methods for predicting protein-RNA interactions could be achieved by integrating the results obtained from scRNA-seq data with complementary information independent of the expression of the RBP and its putative targets. For this reason, we used *cat*RAPID (14, 15) to calculate the interaction propensity of the inferred RBP-RNA pairs and we employed this score to filter out those which are not likely to represent direct interactions (see **Methods** for details).

To find the optimal threshold for the *cat*RAPID score, we tried several cutoff values and recalculated the EPR after removing the inferred interactions whose interaction propensity was lower than the employed cutoff. For most of the datasets and inference algorithms, the EPR increases with the threshold on *cat*RAPID interaction propensity (**Supplementary Figure S6**). We selected 30 as the optimal cutoff for downstream analyses, since for larger values the EPR starts decreasing for some algorithms and datasets. **Figure 2A** shows the EPR obtained after filtering the inferred GRNs with *cat*RAPID for each algorithm and dataset, together with the percentual relative difference with the original EPR value, i.e. the one obtained before *cat*RAPID filtering. Notably, *cat*RAPID leads to an increase of the EPR for all datasets and methods, except for SINCERITIES for the two K562 SCAN-seq2 datasets and DeePSEM for the K562 CEL-seq dataset, providing a relative improvement of the EPR of 17.6%, on average. We notice that ARACNe and TENET_A are among the methods with the largest relative improvement in EPR after filtering based on *cat*RAPID (**Figure 2A** and **Supplementary Figure S7A**), despite both algorithms making use of the DPI to eliminate indirect interactions. DeePSEM and SINCERITIES instead show the least enhancement. Regarding the datasets, the HepG2 Smart-seq2 dataset benefits more than the others from *cat*RAPID, while the K562 CEL-seq and SCAN-seq2 (UMI200) datasets improve less. We also highlight the improvement in the percentage of true positive interactions given by *cat*RAPID (**Supplementary Figures S4 and S5**).

**Figure 2.**
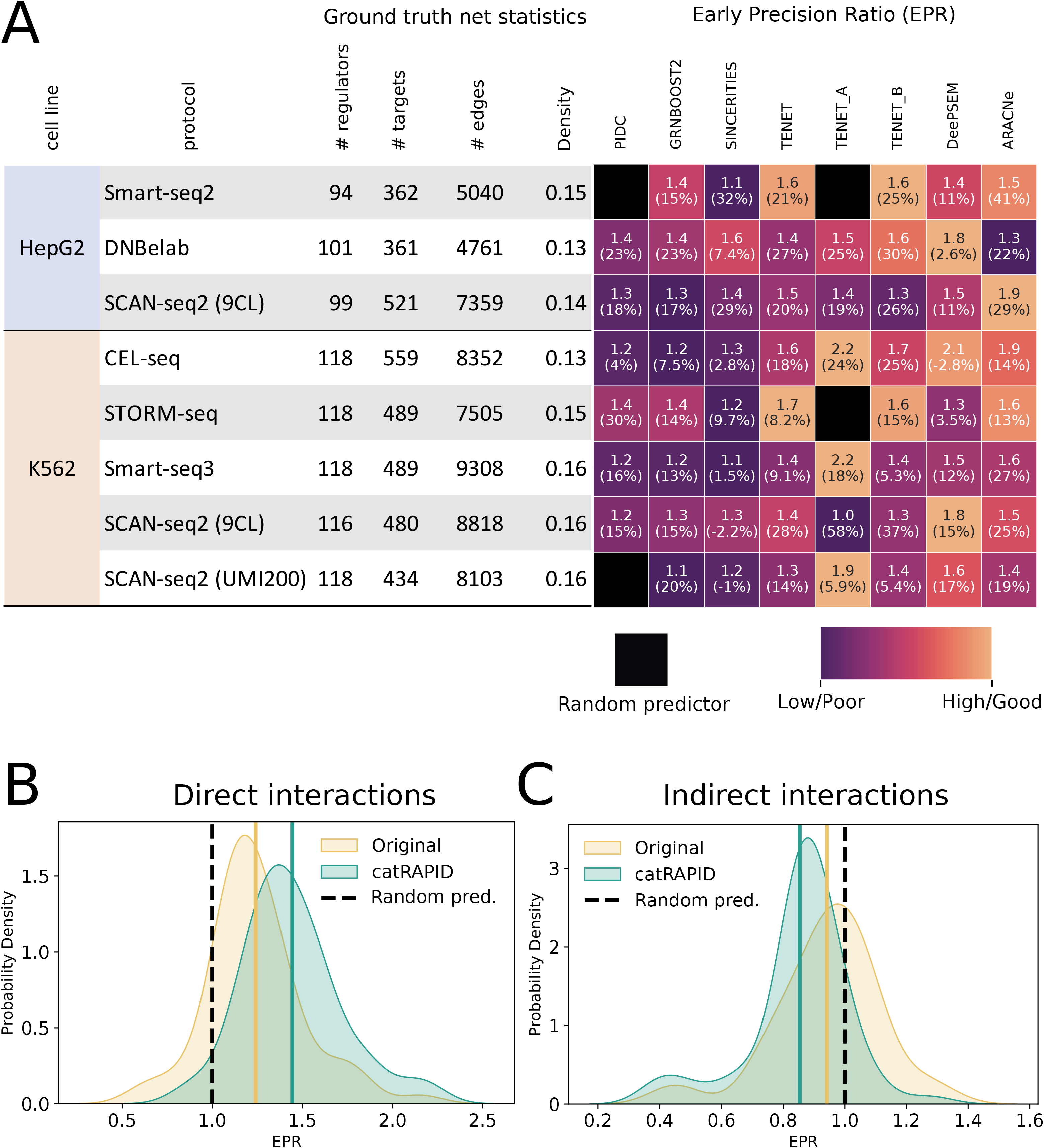
Performances obtained after filtering the inferred RBP-RNA interactions using the *ca*tRAPID algorithm. (**A**) Heatmap showing the EPR measured for each scRNA-seq dataset and GRN inference method after the *cat*RAPID-based filter of the inferred networks. The number in brackets in each cell indicates the relative percentage difference in EPR between the rankings filtered using *cat*RAPID and the original ones. A black box indicates EPR smaller than the one of a random predictor. The colors in the heatmap are scaled between 0 and 1 by row, ignoring values less than that of a random predictor. The table on the left shows the statistics of the eCLIP ground truth networks for each dataset. (**B**) Probability densities of the EPR across methods and datasets for the original rankings and those filtered using *cat*RAPID. The ground truth network is given by eCLIP interactions (p-value= 6.8 x 10^-5^, Kolmogorov-Smirnov test). The black dashed line shows the EPR for a random predictor. (**C**) Same as B, but for indirect RBP-RNA interactions obtained by removing eCLIP interactions from shRNA RNA-seq ones (p-value=7.3 x 10^-4^, Kolmogorov-Smirnov test). In all panels we used scRNA-seq datasets with RBPs included in the eCLIP data and the top 500 HVGs.

Previous work highlighted that the poor performance of inference methods, when evaluated against cell type-specific ChIP-seq data, is due to the presence of indirect interactions, as witnessed by the better performance achieved when using the STRING database as ground truth (31). With the two cell lines under study, we have the possibility of testing this hypothesis for RBP-RNA interactions using shRNA RNA-seq data for the same cell lines, available from the ENCODE project (see **Methods** for details) (4). To evaluate whether inference methods are also predicting indirect interactions, we tested their ability to infer the interactions of RBP with the RNAs deregulated upon their knock-down, from which we removed eCLIP-derived targets to obtain only putative indirect interactions (see **Methods**). While *cat*RAPID causes a shift toward higher EPR values in the case of direct (eCLIP) interactions (**Figure 2B**, Kolmogorov-Smirnov test’s p-value= 6.8 x 10^-5^), for indirect interactions (shRNA RNA-seq) it produces the opposite trend (**Figure 2C**, Kolmogorov-Smirnov test’s p-value= 7.3 x 10^-4^), meaning that it effectively removes indirect interactions. Overall, GRNBOOST2 and ARACNe are the inference methods most prone to detect indirect interactions, while TENET_A infers the least indirect interactions, since it applies the DPI downstream of GRN inference to trim them out (**Supplementary Figures S8 and S9).** Interestingly, also ARACNe applies the DPI but it does not work as effectively as TENET_A, possibly indicating that transfer entropy is more suited than mutual information for the prediction of gene interactions, since the latter does not quantify a directional information flow.

### Predicting protein interactions with long non-coding RNAs

So far we focused on HVGs, which are mostly composed of protein-coding genes. Consequently, the inferred RBP-RNA networks can be confounded by the presence of protein-protein interactions, as mentioned above, while they would not be present when considering only lncRNAs in the datasets. For this reason, for each scRNA-seq experiment, we compared the EPR of the RBP-RNA interactions inferred from two different datasets, built using the top 400 highly variable mRNAs and the top 400 highly variable lncRNAs, respectively. We chose a smaller set of genes compared to the previous analyses since lncRNAs are less represented in the scRNA-seq datasets, due to their smaller absolute number in the transcriptome compared to mRNAs and to their lower expression. In agreement with our expectations, in most of the cases (60%, which becomes 87.5% upon using the *cat*RAPID filter) the performance of the inference algorithms is higher for the lncRNA datasets than for the mRNA ones (**Figure 3 and Supplementary Figure S10**). While for the mRNAs the top performing method is DeePSEM and the best dataset is K562 CEL-seq, as already discussed for the datasets including both types of HVGs, ARACNe emerges as the best method for the inference of RBP-lncRNA interactions. However, we highlight that the smaller number of ground truth eCLIP interactions for lncRNAs might penalize the performance of TENET_A in some datasets, which instead performs very well for the K562 STORM-seq and Smart-seq3 datasets, the newest full length protocols that provide a more precise measurement of lncRNAs expression levels compared to the others. We also evaluated the performances after the *cat*RAPID-based filter (**Figure 3**) and observed that the predictive ability further increases for lncRNAs, even more than for mRNAs. Specifically, the curves of the EPR as a function of the *cat*RAPID interaction propensity threshold (**Supplementary Figure S11**) show that higher values of the EPR are reached for lncRNAs than for mRNAs; the mean EPR over datasets and inference methods is 1.22 for mRNAs and 1.47 for lncRNAs in the original rankings, while it becomes 1.39 for mRNAs and 2.61 for lncRNAs after the *cat*RAPID-based filter with threshold on the interaction propensity set at 30 as in previous analyses. Moreover, for every inference method almost all the datasets with lncRNAs show a monotonic increase of the EPR as a function of the threshold on *cat*RAPID score, especially for the STORM-seq and Smart-seq3 sequencing protocols (**Supplementary Figure S11**).

**Figure 3.**
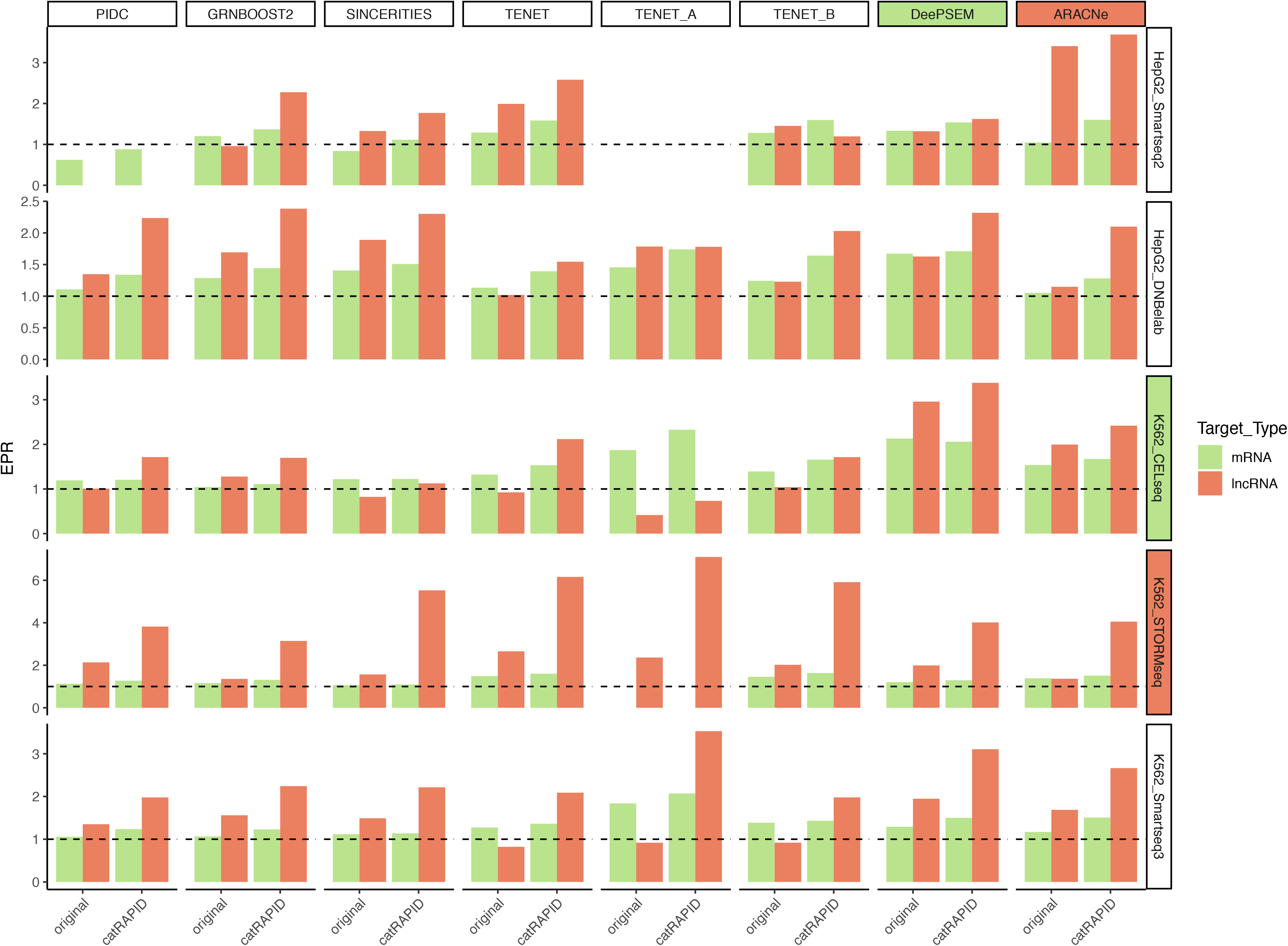
Comparing the inference of RBP-mRNA and RBP-lncRNA interactions. Bar plots showing the EPR measured for datasets with eCLIP RBPs and 400 HVmRNAs or 400 HVlncRNAs, for each GRN inference method (columns) and scRNA-seq dataset (rows). A filled box for an inference method or a dataset indicates that it has the highest EPR (on average), with the color corresponding to mRNAs or lncRNAs, as indicated in the legend. In each bar plot we show the comparison between the performances obtained for mRNA and lncRNAs before (“original”) and after (“*cat*RAPID”) the *cat*RAPID-based filter. The dashed black line indicates the EPR of a random predictor.

Given the large difference in ground truth network size and density between mRNAs and lncRNAs (**Supplementary Figure S10**), for each scRNA-seq dataset we performed 100 random samplings of the eCLIP ground truth networks for mRNAs, to match the statistics obtained for lncRNAs in the same dataset. Next, we compared the EPR values of the lncRNAs with the distribution of EPR values obtained from the samplings of the mRNA ground truth networks, computing an empirical p-value (see **Supplementary Materials**). In **Supplementary Figure S12** we show that, for most of the inference methods and datasets, the EPR values for RBP-lncRNA interactions are significantly higher than those obtained for the downsampled RBP-mRNA interactions. Moreover, *cat*RAPID increases the number of statistically significant comparisons, confirming its usefulness in supporting the prediction of RBP-lncRNAs interactions from scRNA-seq data. Finally, we note that the percentage of true positives is generally smaller in datasets with lncRNAs compared to mRNAs, due to the small size and sparsity of the eCLIP RBP-lncRNA ground truth network, but it gets a large boost after the *cat*RAPID-based filter, especially for the K562 STORM-seq dataset (**Supplementary Figures S13 and S14**).

### Identification of hub genes in protein-RNA networks

One common application of GRN inference from scRNA-seq data is the identification of hub genes (65, 78), intended as genes that regulate a large number of targets, with the goal of discovering novel master regulators (in particular TFs) of the biological process under study. A recent systematic evaluation of GRN inference methods in terms of topological network properties showed that, despite the moderate performance of the algorithms in predicting the correct TF-target edges, the identification of hub genes is generally more reliable (39). Thus, we tested the capability of the methods in predicting hub RBPs and, in addition to previous studies, we also considered “hub RNAs”, defined as RNAs that are regulated by a large number of RBPs. Examples of such RNAs include mRNAs, some of which encode for important regulators of cellular functions like Cyclin D1, c-Fos and Bcl-2, whose 3’UTR contain AU-rich elements recognized by multiple proteins (79), and very long non-coding RNAs, such as MALAT1, NEAT1 and NORAD, whose sequence provides a platform for the binding of multiple factors that are relevant for phase separation (80–82).

Hub RBPs and RNAs were defined according to the out- or in-degree centrality, respectively, computed on the nodes of the ground truth network (see **Supplementary Materials**). Following a previous work (39), we tested the performance of the inference methods using the Jaccard coefficient ratio (JCR), which is the ratio between the Jaccard coefficient of the hubs of the inferred and ground truth networks and the Jaccard coefficient that a random predictor would achieve.

In **Figure 4** we show the results for the datasets that achieve, on average over the inference algorithms, the best performances in terms of JCR in identifying hub RBPs (HepG2 Smart-seq2) and hub mRNAs or lncRNAs (K562 CEL-seq).

**Figure 4.**
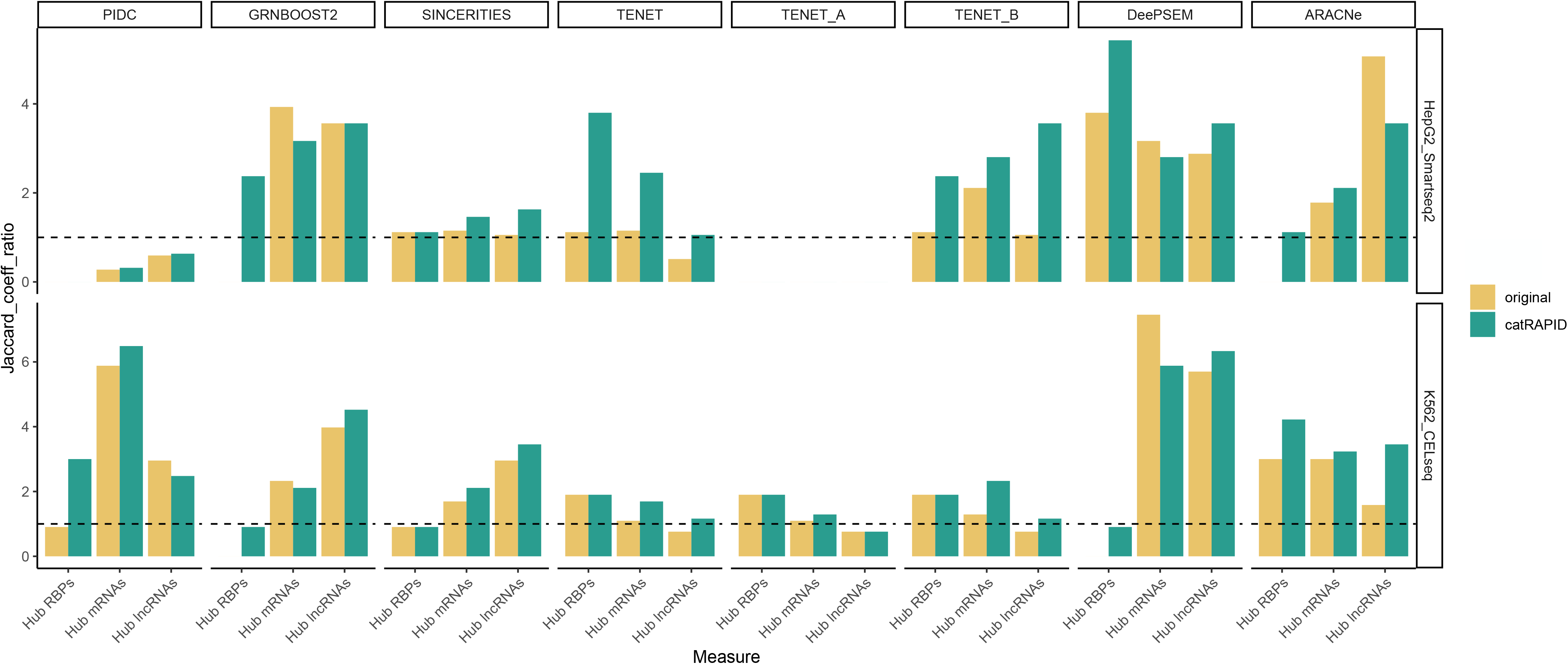
Identification of hub RBPs and hub RNAs in protein-RNA networks. Bar plots showing the JCR for hub RBPs, hub mRNAs and hub lncRNAs before (“original”) and after (“*cat*RAPID”) the *cat*RAPID-based filter, for the HepG2 Smart-seq2 and K562 CEL-seq datasets, that, on average, achieve the best performance in identifying hub RBPs and hub RNAs, respectively. The dashed black line indicates the JCR of a random predictor.

For the prediction of hub RBPs, the filter with *cat*RAPID preserves or increases the value of the JCR in 95% of the cases for the datasets with 500 HVGs and 94% of the cases for the datasets with 1000 HVGs (**Supplementary Figures S15** and **S16**). The K562 SCAN-seq2 (UMI200) and HepG2 SCAN-seq2 (9CL) datasets achieve the best performance before the filter with *cat*RAPID, for datasets with 500 and 1000 HVGs, respectively, while the HepG2 Smart-seq2 dataset benefits most from the filter, becoming the dataset with highest JCR, on average, in both cases (**Figure 4** and **Supplementary Figures S15** and **S16**). Regarding the inference methods, TENET and TENET_B, followed by SINCERITIES and DeePSEM, are the top methods in finding hub RBPs in datasets with 500 HVGs (**Figure 4** and **Supplementary Figure S15**) before the *cat*RAPID-based filter. SINCERITIES becomes the top performing method after the *cat*RAPID filter, followed by DeePSEM and PIDC. For datasets with 1000 HVGs PIDC is the best method, followed by DeePSEM and ARACNe, both before and after *cat*RAPID (**Supplementary Figure S16**).

Regarding the identification of hub mRNAs and lncRNAs, the K562 CEL-seq dataset achieves, on average, the highest JCR, both before and after the *cat*RAPID-based filter, apart from the datasets with 1000 HVGs, in which the top dataset after such filter becomes the K562 SCAN-seq2 (9CL) (**Figure 4** and **Supplementary Figures S15, S16 and S17**). The filter with *cat*RAPID preserves or improves the value of the JCR in 83% of the cases for datasets with 500 HVGs, 75% for datasets with 1000 HVGs and 85% for datasets with 400 lncRNAs. The best method for identifying hub mRNAs is DeePSEM, followed by PIDC, for datasets with 500 HVGs, while it is the opposite for datasets with 1000 HVGs, and this stands both before and after the *cat*RAPID-based filter. Finally, DeePSEM, followed by GRNBOOST2 and ARACNe, is the top performing method in identifying hub lncRNAs.

### Cross-validation of the method using RIP-seq data from murine cells

Subsequently, we assessed the efficacy of the aforementioned methods in predicting RBP-RNA interactions detected through another experimental technique in differentiating cells originating from a distinct organism. To this end, we first considered the C2C12, an immortalized mouse cell line that recapitulates myoblasts to myotube differentiation, for which Split Pool Ligation-based Transcriptome sequencing (SPLiT-seq) scRNA-seq and single-nuclei RNA-seq (snRNA-seq) datasets at 0 (myoblasts) and 72 hours (myotubes) of differentiation are available (83). We inferred the differentiation trajectory from myoblasts to myotubes, selected the top 500 and 1000 HVGs for each dataset and ran the GRN inference (see **Supplementary Materials; Supplementary Figure S18**). We focused the evaluation on the ADAR1 deaminase, which plays a prominent role in skeletal myogenesis, suppressing apoptosis at the myoblast stage and facilitating the myoblast to myotube fate transition (84). ADAR1 target RNAs have been identified in C2C12 cells at 0h and 72h through RNA immunoprecipitation followed by RNA-sequencing (RIP-seq) (84); we highlight that these targets are specific of each time point, thus they represent a restricted list. The results for the myoblasts snRNA-seq dataset are shown in **Figure 5A**. We notice that the performance in terms of EPR is quite robust across algorithms and that *cat*RAPID increases the EPR for all of them except ARACNe, producing, on average, a relative improvement in EPR of 54% for the datasets with 500 HVGs and 24% for those with 1000 HVGs. However, we stress that in this case, in which performances are tested for the interactions of a single RBP, TENET_A, TENET_B and ARACNe are penalized due to the smaller number of interactions that they output compared to the other methods. In **Figure 5** we show only algorithms for which at least two experimental interactions are present in the inferred network, before the *cat*RAPID-based filter.

**Figure 5.**
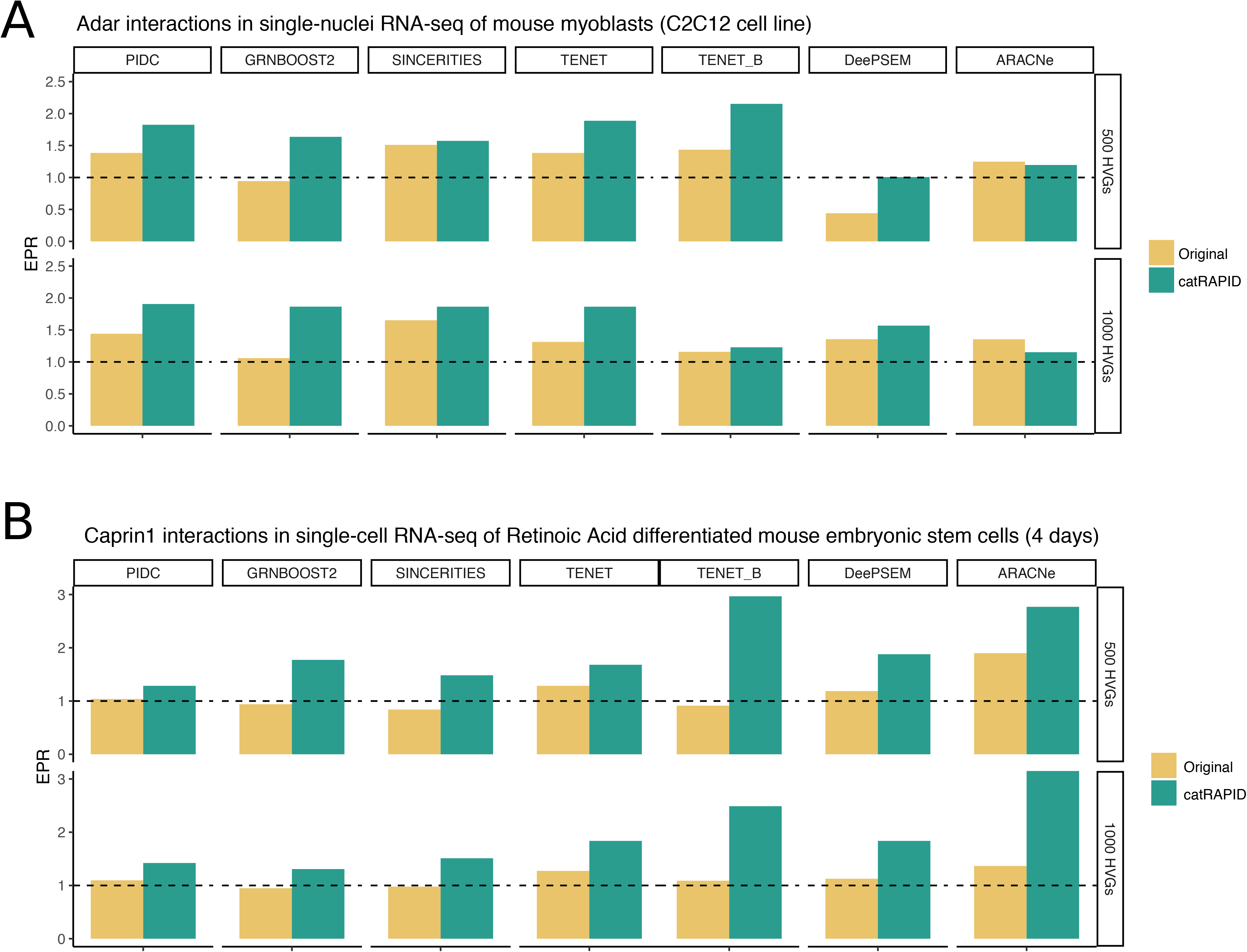
Validation of the method using RIP-seq experiments in mouse cell systems. (**A**) EPR measured for the inference made on the SPLiT-seq dataset of the C2C12 murine cell line recapitulating myoblasts to myotubes differentiation. Performances are tested on the ADAR1 RIP-seq experiments at 0h (myoblasts; here snRNA-seq, see **Supplementary Figure S19A** for scRNA-seq) and 72h (myotubes; see **Supplementary Figure S19B**) of C2C12 differentiation. (**B**) EPR for the SCRB-seq dataset of mouse embryonic stem cells differentiation driven by Retinoic Acid. Performances are tested on the Caprin1 RIP-seq experiments at 0h (undifferentiated mESCs; see **Supplementary Figure S19C**) and 96h (RA-differentiated cells). In both panels we show only algorithms for which at least two experimental interactions are present in the inferred network, before the *cat*RAPID-based filter.

We show the results for a scRNA-seq of myoblasts and snRNA-seq of myotubes, obtained from the same study, in **Supplementary Figures S19A-B**. The results for the myoblasts scRNA-seq (**Supplementary Figure S19A**) are in line with those observed for the snRNA-seq, except for a worse performance of DeePSEM and the increase in EPR after the *cat*RAPID-based filter for ARACNe. Instead, the myotubes dataset (**Supplementary Figure S19B**) shows less consistent results: the filter with *cat*RAPID tends to preserve the EPR values and DeePSEM has higher EPR compared to the other methods, especially for the dataset with 500 HVGs. We hypothesize that the less robust performance obtained for the myotubes dataset is due to the smaller number of ADAR1 targets in myotubes (401 targets vs 3263 targets in myoblasts).

As a second validation system, we considered mouse embryonic stem cells (mESCs) differentiation driven by retinoic acid (RA). We ran the GRN inference on a single-cell RNA barcoding and sequencing (SCRB-seq) dataset in which mESCs were sequenced at 9 time points ranging from 0h to 96h of RA-induced differentiation (85). We evaluated the performance on the RNA targets of the cell-cycle-associated protein 1 (Caprin1), which were obtained at 0h and 96h of RA-induced mESCs differentiation through RIP-seq (86). This protein plays a crucial role during mESC differentiation, regulating an RNA degradation pathway, and its knockout was shown to have a little effect in mESCs while it significantly altered cell differentiation pathways (86). We show the results for the 96h time point in **Figure 5B**. TENET_B and ARACNe outperform the other inference algorithms, especially after filtering the interactions with *cat*RAPID. In this case the EPR of all the methods increases after the *cat*RAPID-based filter. Instead, for the dataset at 0h (**Supplementary Figure S19C**) *cat*RAPID causes a decrease in EPR for some algorithms; this might be explained by the non-essential function of Caprin1 in mESCs (86) and by the smaller number of interactions measured at 0h compared to 96h (1178 Caprin1 targets at 0h vs 2116 at 96h).

### Prediction of RBP co-interactions based on the overlap of inferred targets

The interaction between proteins that bind to common RNA targets can extend beyond their RNA associations and may encompass protein-protein interactions as well (87). Indeed, by binding to shared RNA molecules, the RBPs form a functional partnership that enables coordinated regulation of RNA metabolism and cellular activities (88). The interplay between RBPs at the protein-protein level contributes to the assembly and stabilization of ribonucleoprotein complexes, facilitating RNA processing, transport, and translation (89). These protein-protein interactions can occur through direct physical associations or indirect interactions mediated by bridging factors such as intermediate proteins. Through their interactions, RBPs create dynamic macromolecular complexes that influence RNA localization, stability, and function. Elucidating the protein-protein interactions among RBPs is essential for comprehending the intricate regulatory mechanisms underlying RNA biology and its impact on cellular homeostasis. Therefore, we predicted RBP co-interactions based on their shared RNA targets from the inferred GRNs (see **Methods** for details). To validate the predicted interactions, we considered two cell lines, HEK293T and HCT116, for which scRNA-seq data are available and RBP co-interactions have been experimentally determined through Affinity Purification Mass Spectrometry (AP-MS) and are provided in the BioPlex Interactome database (75).

We considered two scRNA-seq datasets for the HEK293T cell line, obtained with two different protocols (10x and Smart-seq3) (38, 73), and a scRNA-seq dataset from the HCT116 cell line obtained with the Drop-seq protocol (74). For each dataset and inference method, we ranked RBP-RBP pairs based on the fraction of shared targets (see **Methods**). Next, we ran a Gene Set Enrichment Analysis (GSEA) to test the enrichment for experimental RBP co-interactions in the rankings (see **Methods**). In **Figure 6A-C** we show the Normalized Enrichment Score (NES) and the p-value obtained from the GSEA for datasets with 1000, 2000 and 3000 HVGs. We observe that most of the inference methods achieve a significant enrichment, especially for datasets with a higher number of HVGs. In particular, GRNBOOST2 is the best method for the prediction of RBP co-interactions, suggesting that the presence of shared indirect targets can be informative for RBP-RBP interactions, but also DeePSEM, PIDC and ARACNe achieve good performances. The importance of the shared indirect targets for the prediction of RBP-RBP interactions is confirmed by the overall decrease of the NES when the interactions are predicted from the inferred rankings after the *cat*RAPID-based filter (**Supplementary Figure S20**). In **Figure 6D** we show the enrichment plots for the top performing inference methods for each dataset.

**Figure 6.**
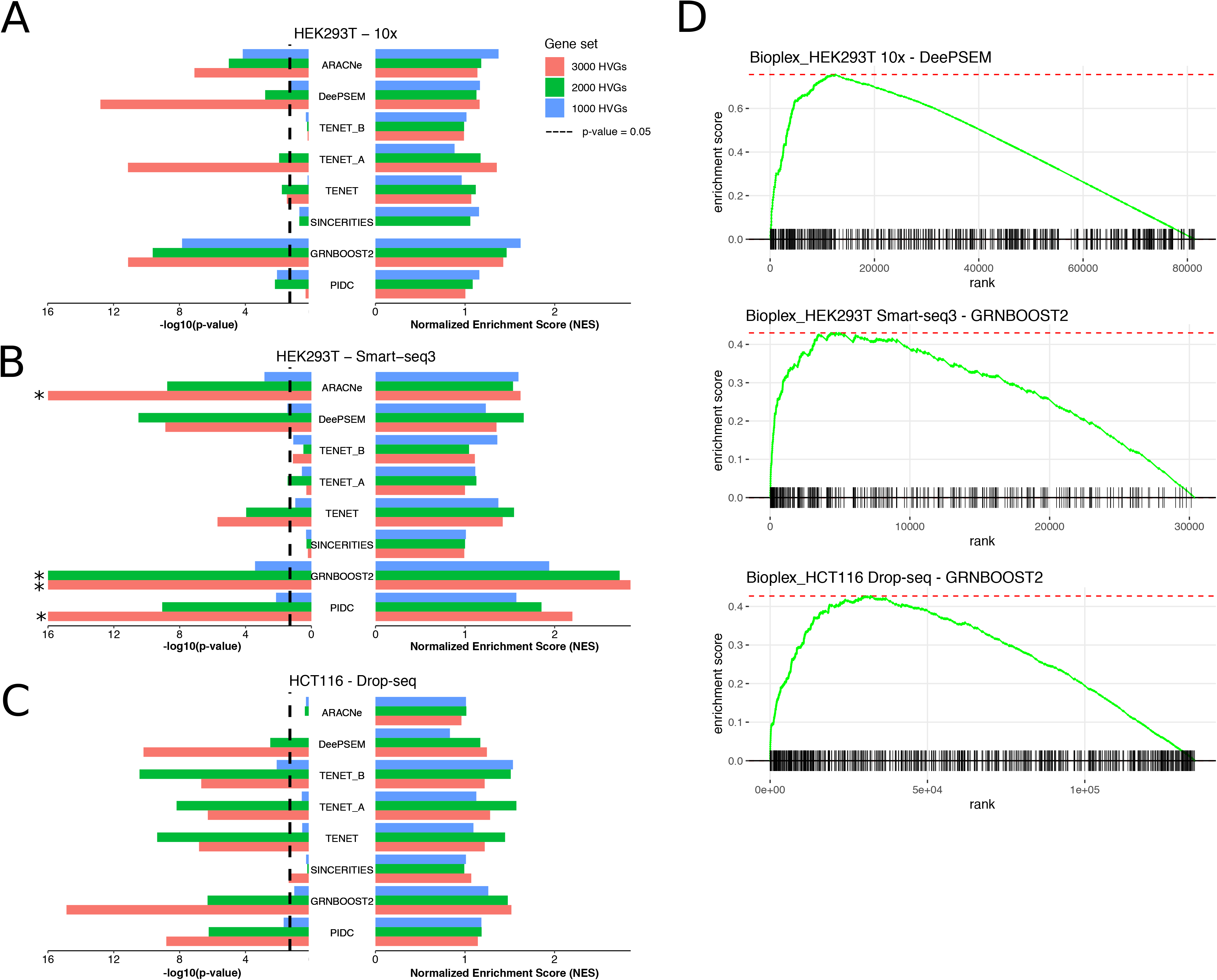
Gene Set Enrichment Analysis of inferred RBP co-interactions. (**A-B-C**) Bar plots showing the - log10(p-value) (left) and Normalized Enrichment Score (NES) obtained from the Gene Set Enrichment Analysis (GSEA) on the inferred RBP-RBP pairs ranked according to the fraction of shared RNA targets, for each GRN inference method. (**A**) scRNA-seq of the HEK293T cell line obtained with the 10x protocol. (**B**) scRNA-seq of the HEK293T cell line obtained with the Smart-seq3 protocol. (**C**) scRNA-seq of the HCT116 cell line obtained with the Drop-seq protocol. p-values capped to 10^-16^ are indicated by a star. (**D**) GSEA enrichment plots for the most significant inference method for each scRNA-seq dataset (3000 HVGs).

## Discussion

In this study, we present scRAPID, a computational pipeline for inferring protein-RNA interactions from single-cell transcriptomic data. We conducted a comprehensive evaluation of the inferred GRNs using various state-of-the-art inference methods across diverse scRNA-seq datasets of different sizes and obtained through different protocols. Importantly, our pipeline is applicable downstream of any inference method, offering flexibility in its usage, and is available at https://github.com/tartaglialabIIT/scRAPID.

We successfully demonstrated the effectiveness of our pipeline in inferring RNA-binding protein (RBP)-RNA interactions. Notably, we achieved similar or even superior performance compared to the inference of TF-target interactions. Furthermore, our observations revealed that the integration of inferred GRNs with *cat*RAPID predictions not only enhanced the inference performance, but also effectively filtered out indirect interactions to a significant extent. Notably, when focusing on RBP-lncRNA interactions, we found even greater improvement, although the task is limited at present by the detection limits of scRNA-seq and eCLIP data used for validation. We highlight that the most recent full-length scRNA-seq protocols, such as Smart-seq3 and STORM-seq, with higher sequencing depth and thus supposed to measure the level of lowly expressed RNAs more precisely, yielded the best results in predicting RBP-lncRNA interactions. The widespread adoption of these protocols, along with improved lncRNA annotation in scRNA-seq (90), is expected to further enhance the prediction of RBP-lncRNA interactions. This development holds great relevance for the identification of functional pathways involving lncRNAs and for the discovery of the underlying mechanisms through which they serve as scaffolds for the formation of protein complexes. By exploring RBP-lncRNA interactions, we could gain insights into the intricate regulatory networks and molecular interactions that contribute to various biological processes. This knowledge is crucial for understanding the roles of lncRNAs and their implications in complex cellular processes, ultimately advancing our comprehension of gene regulation and cellular function.

Moreover, we expanded our investigation beyond binary interactions and demonstrated the ability of the inference methods to predict hub RBPs, hub mRNAs, and lncRNAs. The pipeline’s validation encompassed different organisms and experimental techniques used to obtain protein-RNA interactions. Additionally, we showed the feasibility of predicting RBP-RBP interactions based on their shared targets in the inferred GRNs.

A recent work showed that RBP regulators can be predicted by integrating the GRNs inferred from scRNA-seq data with the information from the RBP binding motifs (36). However, this study uses only GENIE3 for GRN inference, which we did not use since it is very slow and computationally demanding. However, we considered its faster implementation, GRNBOOST2, which has been previously shown to have very similar performance (31), and we found that it shows the lowest performance in predicting RBP-RNA interactions, although it has better performance in detecting hub RBPs. Moreover, the study is limited to the known RBP binding motifs, while our pipeline can be used on any protein, even not necessarily a known RBP.

To better select an appropriate inference method based on the specific task, we conducted a systematic analysis. Our findings indicated that DeePSEM and TENET are the best methods for inferring binary RBP-RNA interactions. DeePSEM was particularly effective for small datasets, while TENET_A was more suitable for larger datasets due to its strict filter on indirect interactions. For inferring RBP-lncRNA interactions, ARACNe and DeePSEM performed well. All inference methods demonstrated proficiency in identifying hub RBPs and RNAs. Notably, PIDC and DeePSEM excelled in hub prediction, while GRNBOOST2 was the top-performing method for predicting RBP co-interactions, despite its tendency to predict more indirect interactions. DeePSEM, PIDC and ARACNe also achieved good performance in this context.

The evaluation of our pipeline on various prediction tasks allowed us to uncover the strengths and weaknesses of different GRN inference algorithms. Methods quantifying the statistical dependence of gene pairs, such as TENET, performed well in inferring binary RBP-RNA interactions but struggled in "global" tasks such as hub detection and RBP co-interactions. In contrast, PIDC, which employs a multivariate information measure between gene triplets, was not effective in identifying RBP-RNA interactions but excelled in hub prediction. Regarding indirect interactions, the application of the DPI effectively filtered them out in TENET_A and TENET_B but not in ARACNe, which was prone to inferring indirect interactions, possibly indicating that transfer entropy should be preferred to mutual information for the prediction of direct regulatory genetic interactions. DeePSEM emerged as the most flexible method, exhibiting good performance in both "local" and "global" prediction tasks. Indeed, its deep neural network based on the Structural Equation Model (SEM) learns features from the scRNA-seq data that enable data embedding, simulation and GRN inference with the same model.

The ability to predict common targets of RBPs is fundamental to identify elements that co-assemble in phase-separated assemblies such as SGs (91). Indeed, RNA is a key component of SGs and it has been proposed that rising levels of ribosome-free mRNAs drive SG formation during stress (92). The molecular composition and the function of proteins in the compartmentalization and the dynamics of assembly and disassembly of phase-separated assemblies is being studied in detail, but the role of RNA in these structures still remains largely unknown (93). RNA can function as molecular scaffolds recruiting multivalent RBPs and their interactors to form higher-order structures (93). Following our approach, we showed that beyond predicting the RNA interactors of proteins that mediate SG condensation, like Caprin1, common RNA targets of RBPs can be inferred to a remarkable extent, which could be in the future exploited to identify transcripts favoring the assembly of protein complexes and their phase separation.

The combination of the predictions of binary RBP-RNA interactions with RBP co-interactions might lead to the development of methods for predicting cell-type specific ribonucleoprotein complexes. By integrating the cell-type resolution of GRN inference from single-cell transcriptomic data with the structural and physico-chemical information encoded by *cat*RAPID, our pipeline enables cell-type-specific prediction of new protein-RNA interactions — an exceptionally challenging task from an experimental point of view (6).

A promising future application of our strategy consists in the integration of other single-cell omics data to predict GRNs at multiple layers of gene expression regulation. In parallel, the advancement in the resolution and throughput of experimental techniques for the detection of protein-RNA interactions will provide more accurate data to test the computational methods. Indeed, massively multiplexed methods for the simultaneous measurement of protein-RNA interactions from tens to hundreds of RBPs, such as antibody barcode eCLIP (94) and Split and Pool Identification of RBP targets (SPIDR) (95), have been recently developed, and they are expected to produce massive interaction datasets in the near future (10).

## Acknowledgments

The authors would like to thank the ‘RNA initiative’ at IIT and all the members of Tartaglia’s lab at CRG, Sapienza and IIT. Especially, we thank Michele Monti and Giorgio Bini for useful discussions.

## Data and code availability

The code to reproduce the analysis and figures in the manuscript is provided at the Github repository https://github.com/tartaglialabIIT/scRAPID. We also include a tutorial to run scRAPID on GRNs inferred on new single-cell transcriptomic datasets, for which the ground truth is not known. The code for GRN inference, evaluation of the performance and plotting is compatible with BEELINE (31) and STREAMLINE (39).

The scRNA-seq datasets used in this study are available from public repositories, listed in **Supplementary Table S1** and discussed in depth in the **Methods** section.

The eCLIP, ChIP-seq and shRNA RNA-seq data for the HepG2 and K562 cell lines are publicly available in the ENCODE project portal (https://www.encodeproject.org/) (42); we report all the eCLIP and ChIP-seq datasets used in this study in **Supplementary Table S2**. Refer to the **Methods** section for their processing.

The RIP-seq data for ADAR1 in C2C12 cells and for Caprin1 in RA-differentiated mESCs used for method validation are available as supplementary tables from the studies (84) and (86), respectively.

We provide a database with *cat*RAPID scores for human and mouse RBP-RNA interactions that can be queried via “curl”; see the Github repository https://github.com/tartaglialabIIT/scRAPID for details and examples queries.

## Funding

Our research was supported by the ERC ASTRA_855923 (GGT), H2020 projects IASIS_727658 and INFORE_825080 and IVBM4PAP_101098989 (GGT) as well as PNRR ‘National Center for Gene Therapy and Drugs based on RNA Technology’’ (GGT).

## Notes

### Competing Interest Statement

The authors have declared no competing interest.

