## Supplementary Materials and Figures for "Prediction of protein-RNA interactions from single-cell transcriptomic data"

#### Quantities used for evaluation

In all the evaluation analyses, we intersected the ground truth networks with the single-cell RNA-sequencing (scRNA-seq) dataset under study. For the RBP-lncRNA interactions, we also removed RBP-RBP interactions from the eCLIP network. Next, the possible edges were computed from the ground truth network considering all the possible combinations of the set of regulators (TFs or RBPs) with the set of possible target genes, which are all the genes selected for the inference, except for the lncRNA datasets, for which the mRNAs corresponding to RBPs were excluded. The rankings inferred by each algorithm were then intersected with the set of possible edges, and self-loops were excluded.
We ranked the inferred networks by descending edge weight as returned by each algorithm. For ARACNe-AP the FDR of the inferred edges is returned, thus we used -log10(FDR) as the edge weight.
Our evaluation pipeline is based on BEELINE, but we provide several extensions to it, written in a compatible way, such as the computation of the percentage of true positive edges in the inferred rankings, the identification of hub RBPs and hub RNAs (which is based on STREAMLINE (39)), the calculation of the Early Precision Ratio restricted to RBP-RNA interactions, the downsampling analysis of RBP-mRNA networks and the prediction of RBP co-interactions. The extensions are provided in our Github repository https://github.com/tartaglialabIIT/scRAPID.

#### Early Precision

The Early Precision (EP) is defined as the fraction of true positives in the top k-edges of the inferred network, where k is the number of edges in the ground truth network. If an inference method returns n < k edges, we consider all the n edges. We adopt this measure following BEELINE (31), since the most confident predicted interactions are expected to be the most interesting for experimental scRNA-seq datasets.

The Early Precision Ratio (EPR) is defined as EPR=EP/d, where d is the expected EP of a random predictor for a given ground truth network, and it equals the ground truth network density, defined as the ratio between the number of true edges and the number of possible ones.

#### True positives in ranking

We include an additional measure of the performance of the inference methods computing the percentage of true positive edges in the ranking of inferred interactions divided in sets of 50 elements. To plot the results as smooth curves, we fit a cubic smoothing spline using the R function “smooth.spline”, with smoothing parameter “spar=0.7”. The corresponding percentage of true positives for a random predictor is given by the ground truth network density multiplied by 100.

#### Hub identification

We used the degree centrality as a measure of hub gene identification (39). Since we deal with directed RBP-RNA networks, we defined hub RBPs based on the out-degree centrality, i.e. the fraction of edges going out from the node of interest, and hub RNAs based on the in-degree centrality, i.e. the fraction of edges going in the node of interest, for hub RNAs. We ranked the ground truth and inferred network nodes (RBPs or RNAs) according to the out or in-degree centrality. We defined as hubs the top 10% of nodes in the ground truth network, indicated as $\Omega_{GT}$ according to the selected measure and we computed the Jaccard coefficient with the same number of nodes obtained from the inferred network, indicated as $\Omega_{INF}$, defined as:

$J(\Omega_{GT}, \Omega_{INF}) = \frac{|\Omega_{GT} \cap \Omega_{INF}|}{|\Omega_{GT} \cup \Omega_{INF}|}$ .

The Jaccard coefficient ratio is the fraction between $J(\Omega_{GT}, \Omega_{INF})$ and the expected value for a random predictor, for which an analytical formula exists (69):

$J(\Omega_{GT}, \Omega_{RAND}) = \frac{p^2}{2p - p^2}$ ,

where $\Omega_{RAND}$ is a set of randomly predicted hubs and $p \in (0, 1]$ is the fraction of nodes in the network defined as hubs (0.1 in our case).

#### Downsampling of eCLIP network for mRNA-lncRNA comparison

To confirm that the performance in terms of EPR for RBP-lncRNA eCLIP interactions compared to RBP-mRNA is not biased by the different size and density of the ground truth networks, we performed a downsampling analysis of the RBP-mRNA networks.
First, we removed RBP-RBP interactions from the eCLIP ground truth. Next, for each scRNA-seq dataset, to obtain a RBP-mRNA network with comparable number of RBPs, RNAs, total number of edges and network density we devised a recursive approach:

- Randomly sample N RBPs from the set of M RBPs in the RBP-mRNA eCLIP network, where N is the number of RBPs in the RBP-lncRNA network;
- If M < N we consider all the M RBPs and we balance the network densities by sampling more targets from the RBP-mRNA network;
- Loop over the selected RBPs and randomly sample one RBP-mRNA interaction at a time;
- Control that the numbers of mRNAs and edges do not exceed those of the corresponding RBP-lncRNA network;
- Remove the sampled interaction from the set of possible ones.

Repeating this procedure 100 times, we obtained RBP-mRNA networks that, on average, have exactly the same features of the corresponding RBP-lncRNA networks. For each RBP-mRNA ground truth network we computed the EPR, for each scRNA-seq dataset and inference method.
From the distribution of EPR values for the downsampled RBP-mRNA networks we computed an empirical cumulative distribution function (ECDF). Finally, the probability to obtain a EPR larger than the EPR for the corresponding RBP-lncRNA ground truth network, indicated as $EPR_{lncRNA}$, was computed from the ECDF in the following way:
$p(EPR > EPR_{lncRNA}) = 1 - ECDF(EPR_{lncRNA})$.
The results are presented in ‘Predicting protein interactions with long non-coding RNAs’ and **Supplementary Figure S12** for each scRNA-seq dataset and GRN inference method, where a star indicates that $p(EPR > EPR_{lncRNA}) < 0.05$, namely that $EPR_{lncRNA}$ is significantly larger than the EPR for RBP-mRNA interactions. The code for reproducing the downsampling analysis is provided in our Github repository https://github.com/tartaglialabIIT/scRAPID.

### Datasets used for validation

#### Differentiating mouse myoblasts dataset

We used a SPLiT-seq dataset applied to the C2C12 myogenic system (70), recapitulating myoblasts to myotube differentiation. The dataset includes scRNA-seq of myoblasts (0h of differentiation), single-nuclei RNA-seq of myoblasts (0h of differentiation) and single-nuclei RNA-seq of myotubes (72h of differentiation) and it is available on GEO with accession number GSE168776. In our analyses we used the 9000-cell library of the short-read SPLiT-seq dataset, selecting the batch C since it has the highest median UMI count and gene count per cell. It includes 1305 single myoblast cells, 1718 single myoblast nuclei and 3288 single myotube nuclei. We computed a diffusion map as described above and we choose as root cell for the computation of the diffusion pseudotime the myoblast cell with the largest value of Diffusion Component 1 in the diffusion map (see section 'Cross-validation of the method using RIP-Seq data from murine cells' and associated **Supplementary Figures S18**). Next, we separated the three cell populations (single-cell myoblasts, single-nuclei myoblasts and single-nuclei myotubes) and we performed selection of the top 500 and 1000 HVGs prior to GRN inference in each population independently.

We used the mouse RBPs included in the RBP2GO database (7), with RBP2GO score larger than 10, and we added to each dataset the RBPs that belong to the full set of the HVGs (i.e. genes that are statistically significant according to the statistical test performed using the Scanpy function "scanpy.pp.highly_variable_genes", without setting the parameter "n_top_genes"). We manually added ADAR1 to the gene set for GRN inference for the datasets in which it does not belong to the set of HVGs. GRN inference on the processed datasets was run as described above.

Next, we obtained RBP-RNA interactions for ADAR1 from a native RNA immunoprecipitation (RIP) combined with RNA-Seq (RIP-Seq) experiment presented in (71), which has been performed in the same cell line (C2C12) and corresponding time points (0h and 72h) as the scRNA-seq dataset. In this study, 3263 and 401 ADAR1 targets were found at 0h and 72 h, respectively. We highlight that the targets were defined based on the log2 fold change of the binding enrichment between the two time points, hence they represent specific targets for the given time point.

We evaluated the performance of each GRN inference method in predicting ADAR1 RIP-seq interactions before and after the filter of the rankings using *cat*RAPID with the same pipeline and evaluation metrics described above.

#### Mouse embryonic stem cells differentiation

We used a SCRB-seq scRNA-seq dataset of mouse embryonic stem cells (mESCs) differentiation driven by retinoic acid (RA) (72). mESCs were sequenced at 9 different time points (0h, 6h, 12h, 24h, 36h, 48h, 60h, 72h, 96h). The UMI count matrices for all the time points are available on GEO with accession number GSE79578. In this case the authors provided all the cells, including bad quality ones, thus we removed cells with less than 2000 total UMI counts as done in the original publication. Then, we computed a diffusion map as described above, which showed a clear trajectory starting from the undifferentiated mESCs, and we selected as root cell for the computation of the diffusion pseudotime the cell at 0h with the smallest value of diffusion component 1 (see section 'Cross-validation of the method using RIP-Seq data from murine cells' and **Supplementary Figure S18**).

Next, we retained the 282 undifferentiated mESCs (0h) and the 137 ones obtained after 4 days of RA-induced differentiation (96h), and we performed the gene selection step for GRN inference as described in the previous paragraph.

The interactions for Caprin1 were obtained from a RIP-seq experiment performed in undifferentiated mESCs and RA-differentiated cells at 96h (73), matching the time points of the scRNA-seq dataset. From the RIP-seq, the authors identified 1178 and 2116 Caprin1 RNA targets at 0h and 96h, respectively.

The evaluation of the inference performance before and after *cat*RAPID-based filter was performed as described in the previous sections.

### Supplementary Figures

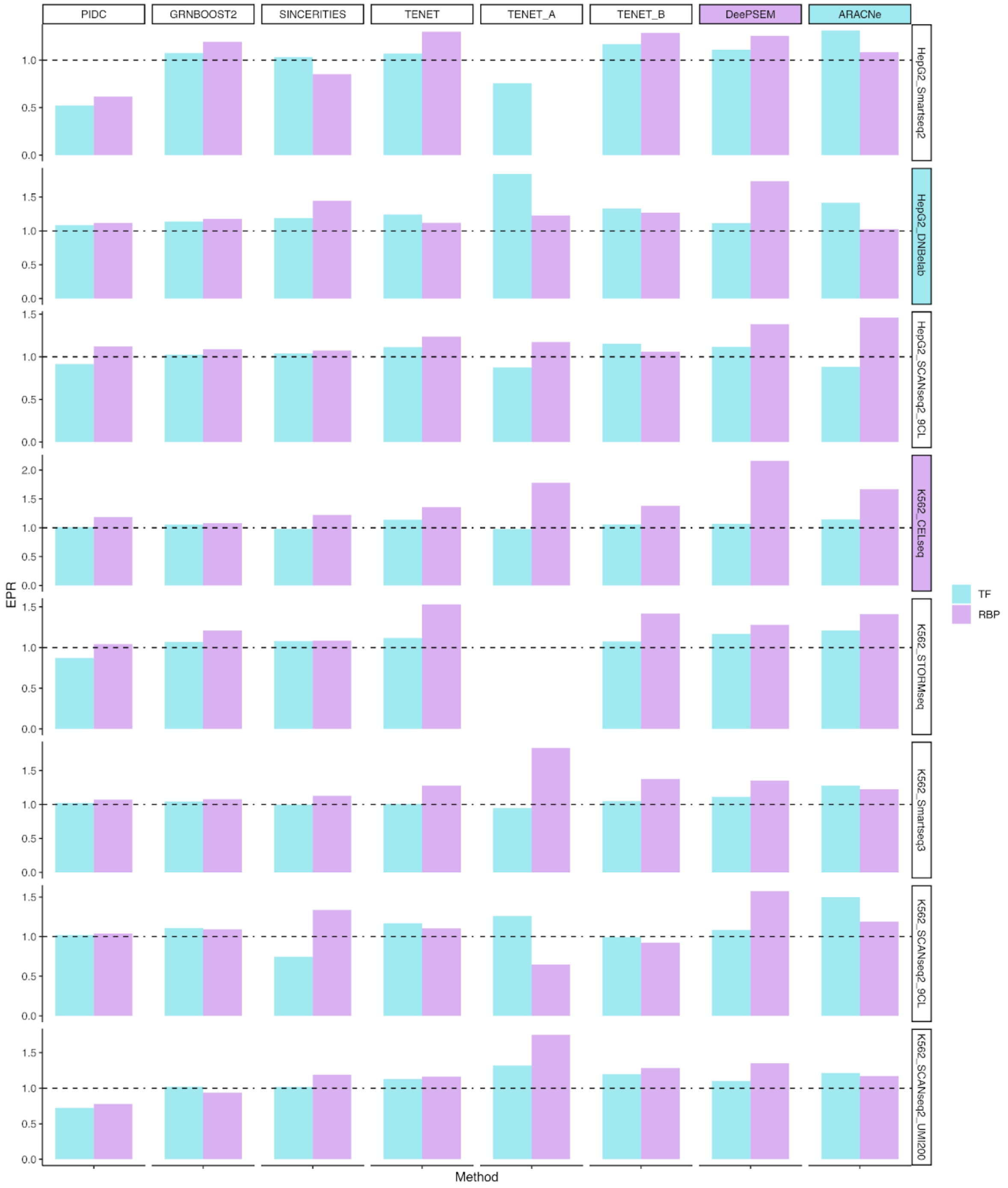


**Supplementary Figure S1**. **Related to Figure 1.** Bar plots showing the Early Precision Ratio (EPR) obtained for TF-target and RBP-target interactions by each GRN inference method (columns), for each scRNA-seq dataset (rows) with 500 HVGs selected. The black dashed line shows the EPR of a random predictor.

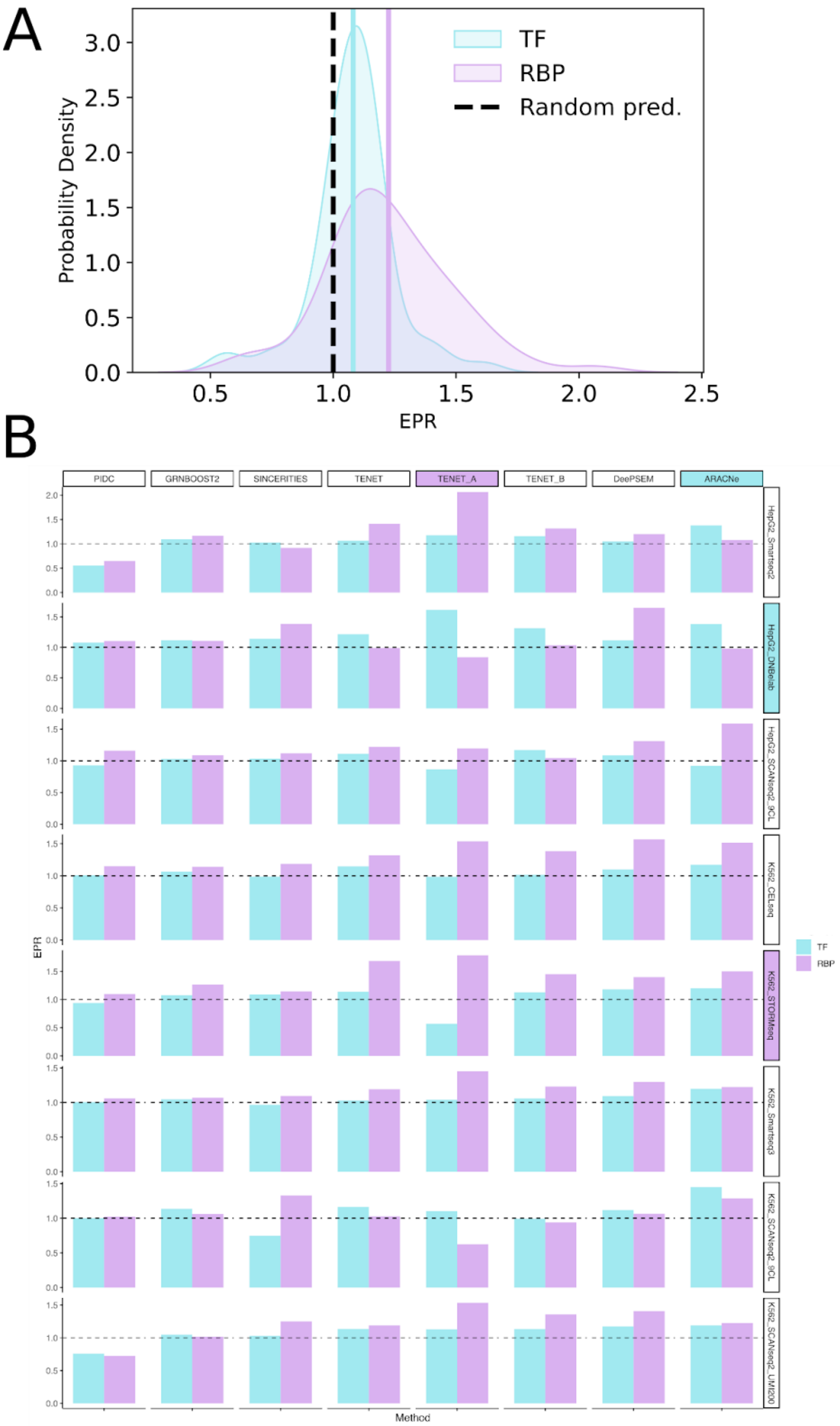
A
B
Probability Density
EPR
TF
RBP
Random pred.
PIDC
GRNBOOST2
SINCERITIES
TENET
TENET_A
TENET_B
DeePSEM
ARACNe
HepG2_Smartseq2
HepG2_DNBelab
HepG2_SCANseq2_9CL
K562_CELseq
K562_STORMseq
K562_Smartseq3
K562_SCANseq2_9CL
K562_SCANseq2_UMI200
Method

**Supplementary Figure S2. Related to Figure 1.** (**A**) Probability density of the EPR across methods and datasets for the TF-target and RBP-target datasets. The ground truth network is given by cell-type specific ChIP-seq and eCLIP interactions for TF-target and RBP-target interactions, respectively (p-value=$4.1 \times 10^{-5}$,Kolmogorov-Smirnov test). (**B**) Bar plots showing the EPR obtained for TF-target and RBP-target interactions by each GRN inference method (columns), for each scRNA-seq dataset (rows). Both panels refer to the analysis with 1000 HVGs selected. The black dashed line shows the EPR of a random predictor.

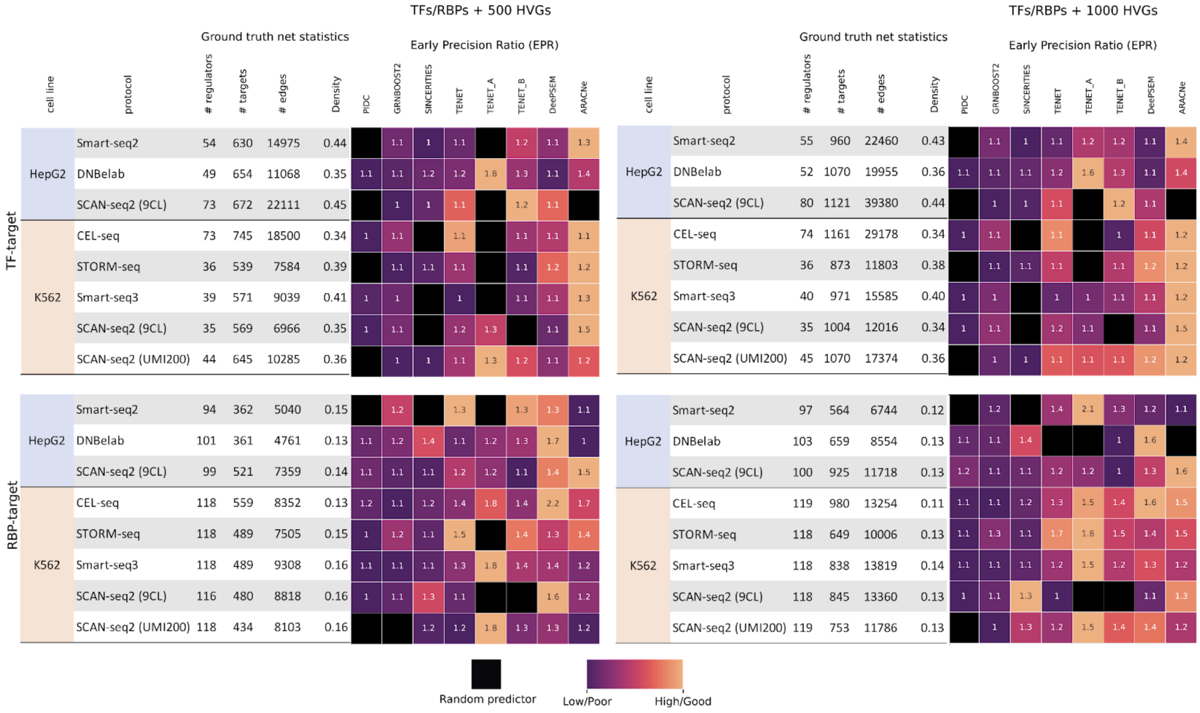


**Supplementary Figure S3. Related to Figure 1.** Heatmaps showing the EPR for each scRNA-seq dataset (rows) and GRN inference method (columns) for the inference of TF-target (top) and RBP-target (bottom) interactions, for datasets with 500 (left) or 1000 (right) HVGs. A black box indicates EPR smaller than the one of a random predictor. The colors in the heatmap are scaled between 0 and 1 by row, ignoring values less than that of a random predictor. The table on the left of each heatmap shows the statistics of the cell-type specific ChIP-seq or eCLIP ground truth network for each dataset.

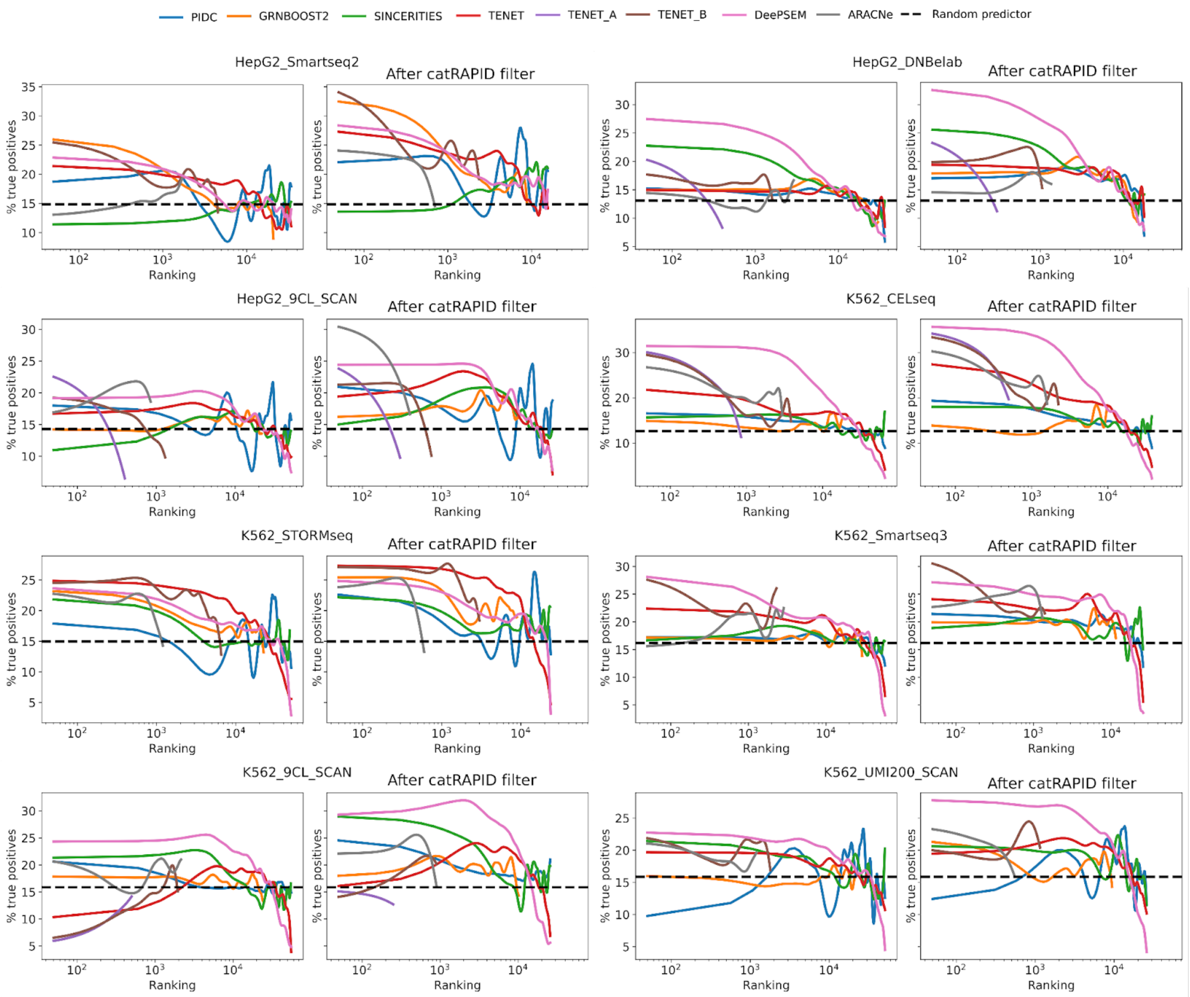


**Supplementary Figure S4**. **Related to Figures 1 and 2.** Percentage of true positive RBP-target interactions, according to the cell-type specific eCLIP ground truth network, as a function of the rank of the interactions inferred from scRNA-seq data, for each algorithm, before and after the *cat*RAPID-based filter, for the datasets with 500 HVGs.

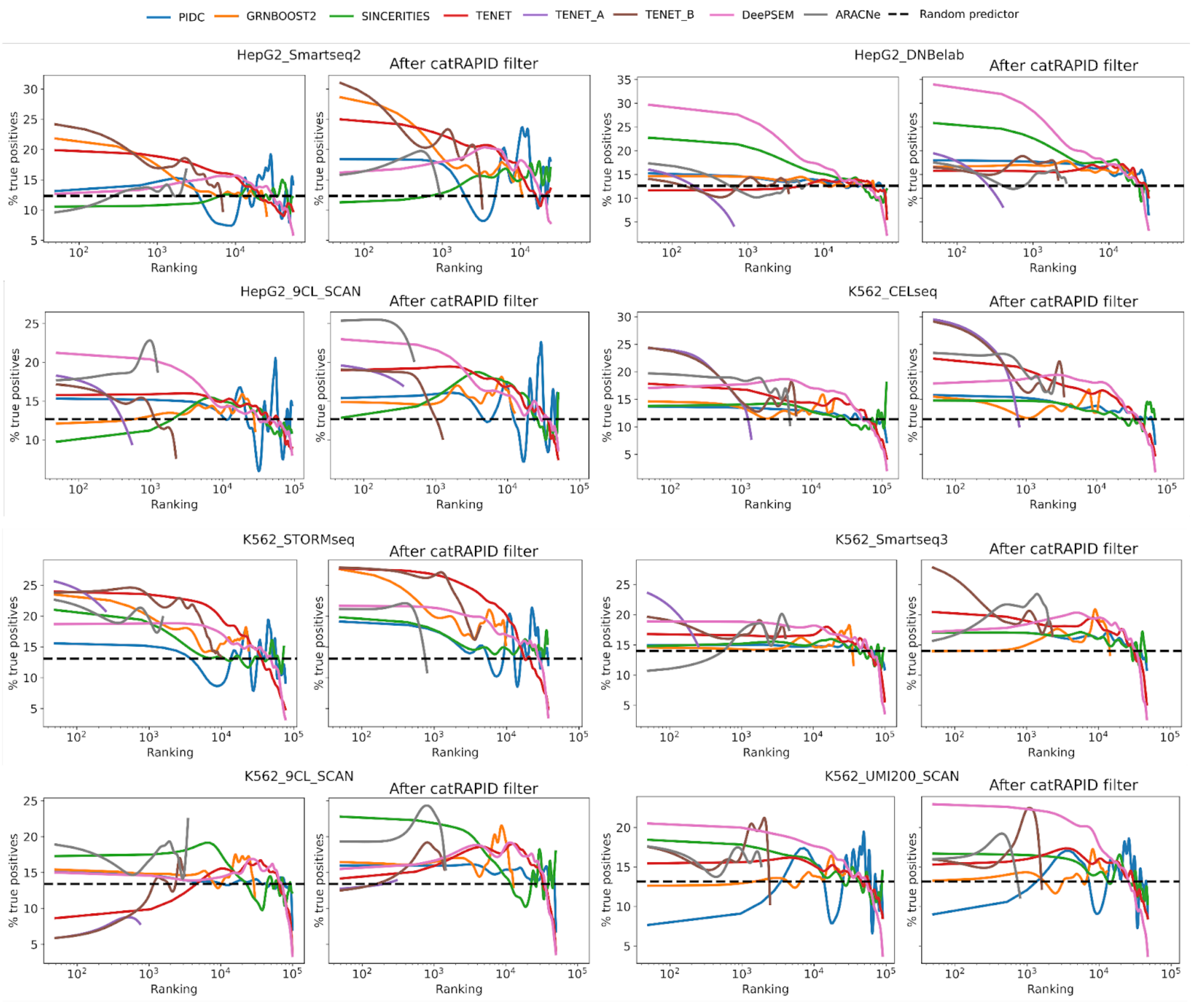


**Supplementary Figure S5. Related to Figures 1 and 2.** Percentage of true positive RBP-target interactions, according to the cell-type specific eCLIP ground truth network, as a function of the rank of the interactions inferred from scRNA-seq data, for each algorithm, before and after the *cat*RAPID-based filter, for the datasets with 1000 HVGs.

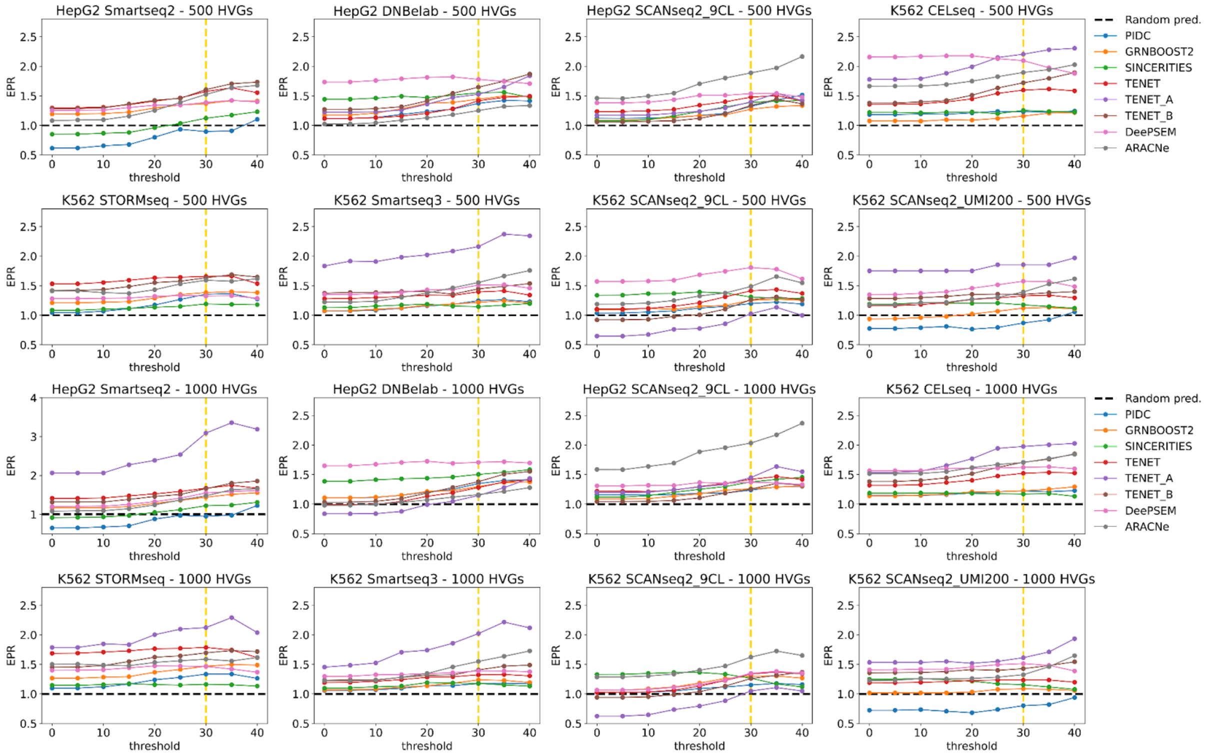


**Supplementary Figure S6**. **Related to Figure 2.** EPR as a function of the threshold on *cat*RAPID interaction propensity used for filtering the GRNs inferred by each algorithm. We show the datasets with 500 HVGs selected in the top two rows and with 1000 HVGs in the bottom two rows. The vertical yellow dashed line indicates the threshold selected for all the analyses. The horizontal black dashed line indicates the EPR of a random predictor. The y-axis range is the same for all panels, except for the HepG2 Smart-seq2 dataset with 1000 HVGs. The ground truth network is given by eCLIP interactions.

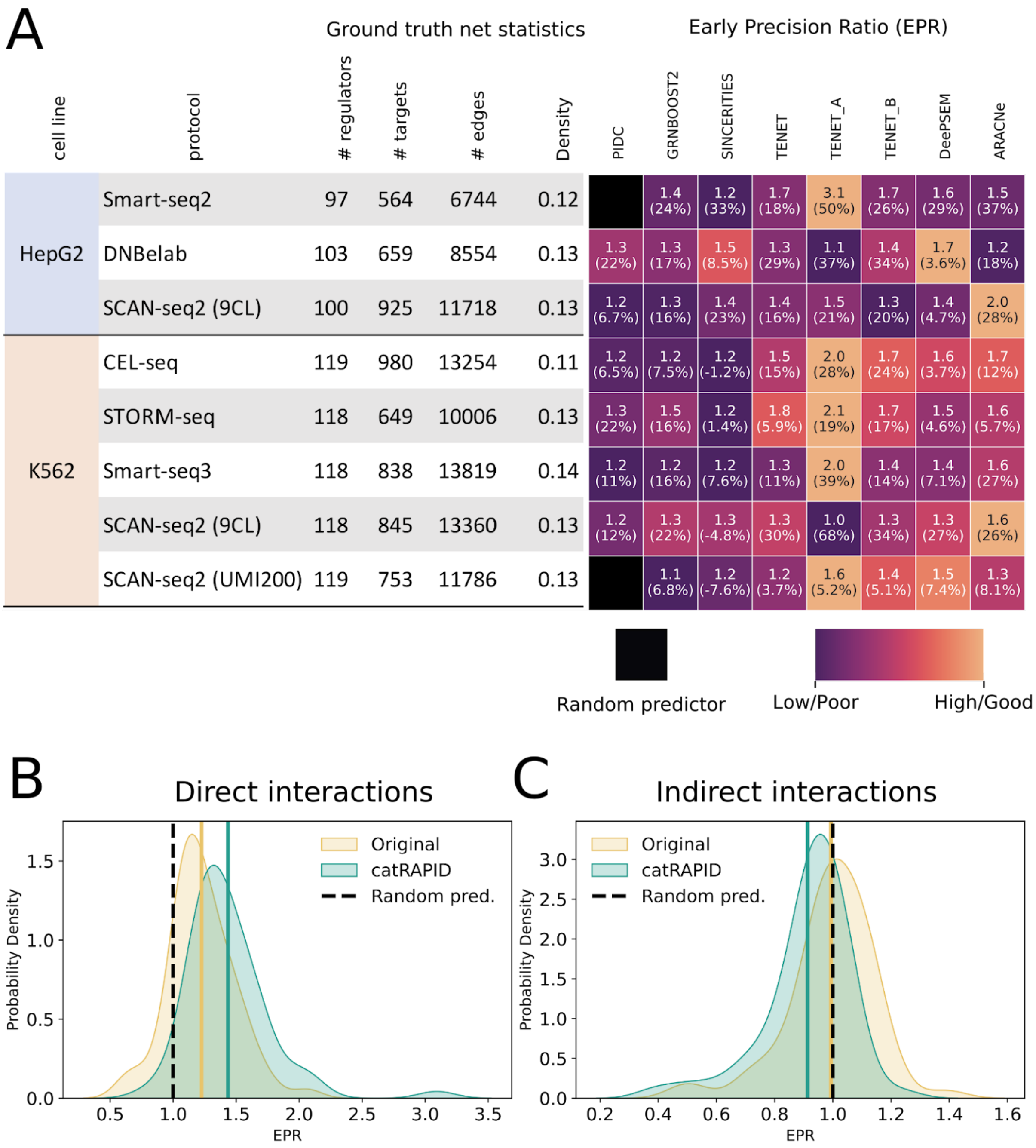


**Supplementary Figure S7. Related to Figure 2.** (**A**) Heatmap showing the EPR for each scRNA-seq dataset and GRN inference method after the *cat*RAPID-based filter to the inferred networks. The number in brackets in each cell indicates the relative difference in EPR between the rankings filtered using *cat*RAPID and the original ones. A black box indicates EPR smaller than the one of a random predictor. The colors in the heatmap are normalized by row. The table on the left shows the statistics of the eCLIP ground truth networks for each dataset. (**B**) Probability densities of the EPR across methods and datasets for the original rankings and those filtered using *cat*RAPID. The ground truth network is given by eCLIP interactions (p-value=4.6 x $10^{-4}$, Kolmogorov-Smirnov test). The black dashed line shows the EPR for a random predictor. (**C**) Same as B, but for indirect RBP-RNA interactions obtained by removing eCLIP interactions from shRNA RNA-seq ones (p-value=p-value=1.9 x $10^{-3}$, Kolmogorov-Smirnov test). In all panels we used scRNA-seq datasets with RBPs included in the eCLIP data and the top 1000 HVGs.

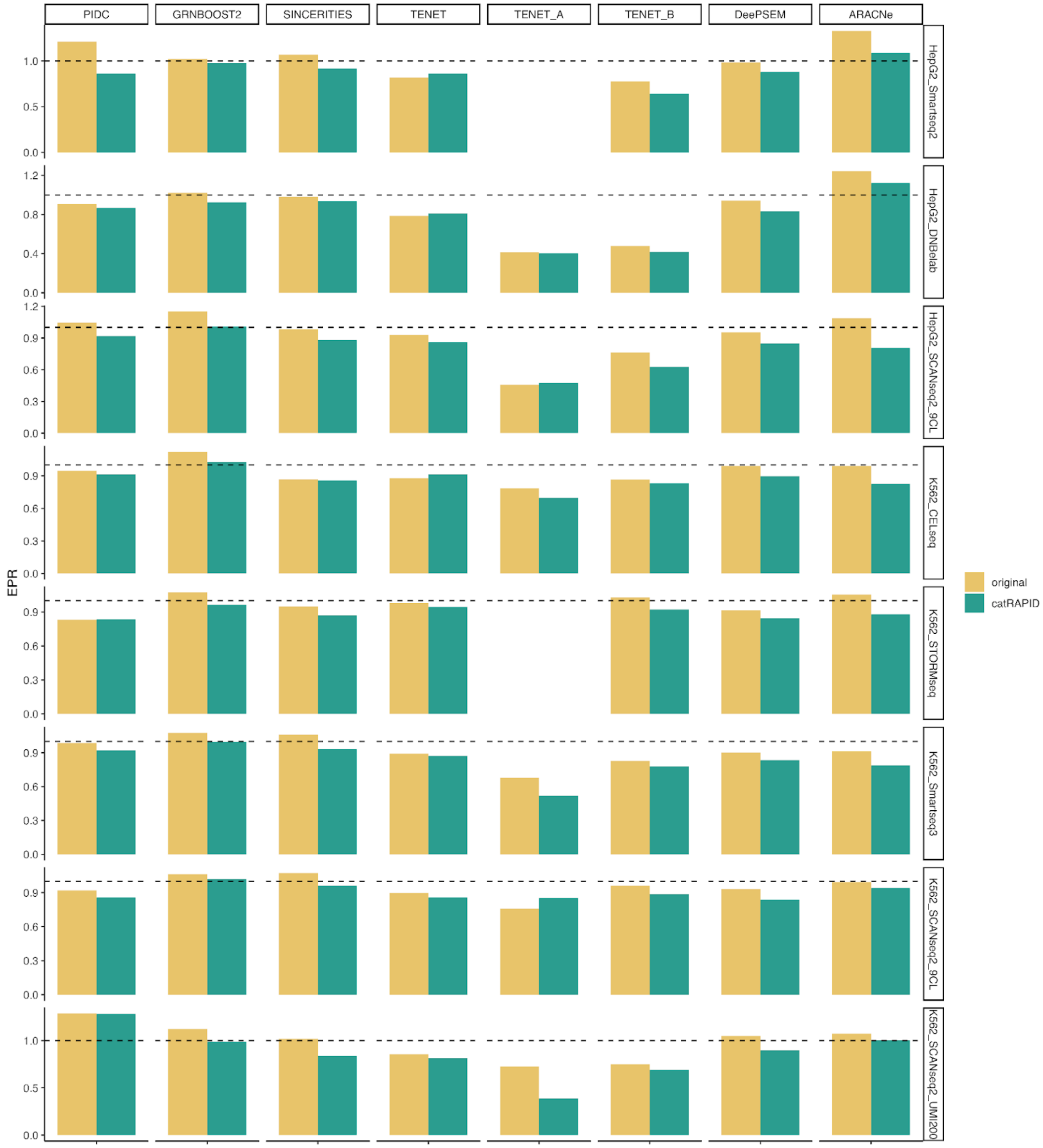


**Supplementary Figure S8. Related to Figure 2.** Bar plots showing the EPR obtained for RBP-target indirect interactions by each GRN inference method (columns), for each scRNA-seq dataset (rows) with 500 HVGs. The black dashed line shows the EPR of a random predictor. The ground truth network is obtained by considering the RBPs present in both shRNA RNA-Seq and eCLIP data, and removing eCLIP interactions from shRNA RNA-Seq ones, thus obtaining indirect interactions.

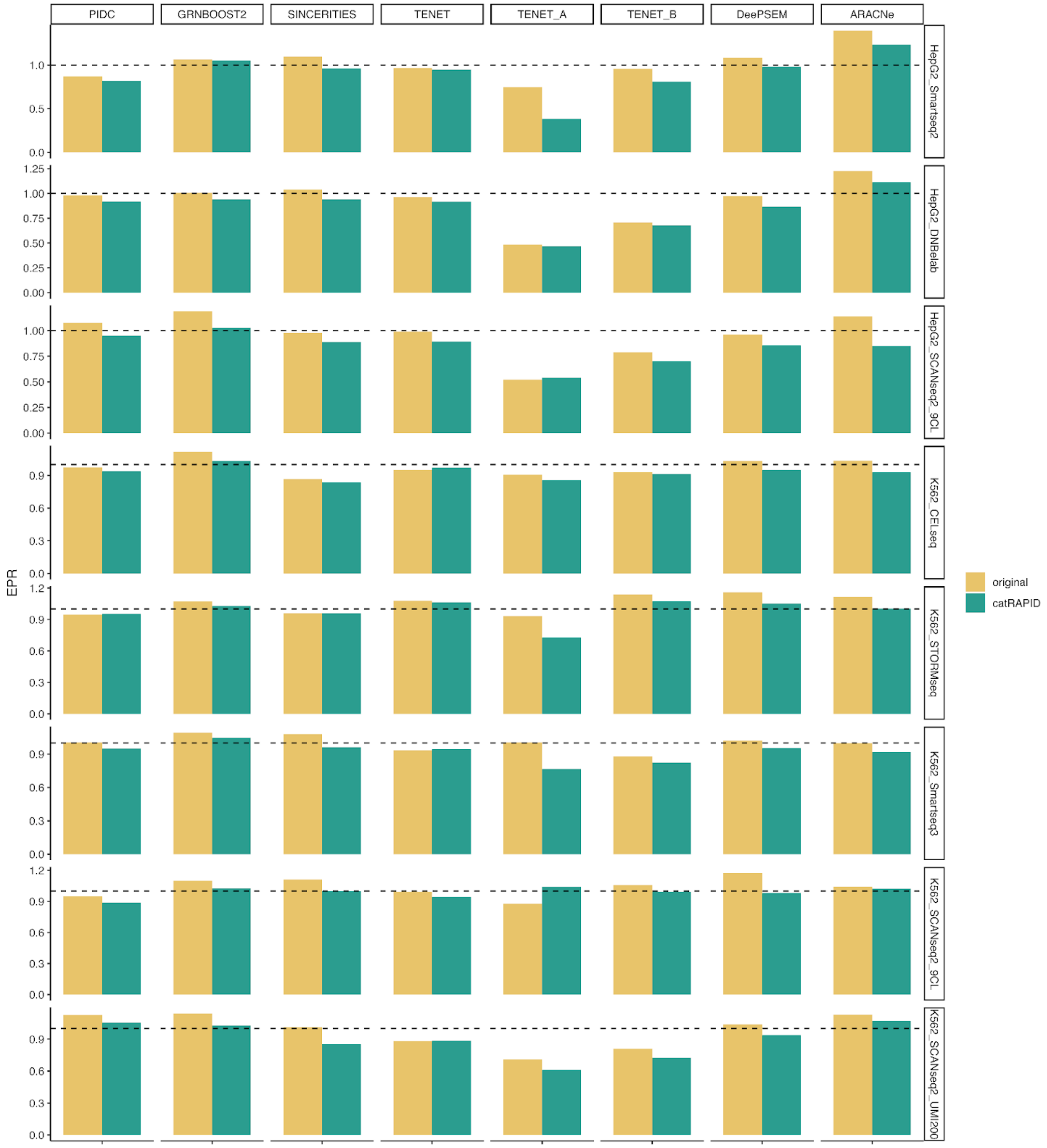


**Supplementary Figure S9. Related to Figure 2.** Bar plots showing the EPR obtained for RBP-target indirect interactions by each GRN inference method (columns), for each scRNA-seq dataset (rows) with 1000 HVGs. The black dashed line shows the EPR of a random predictor. The ground truth network is obtained by considering the RBPs present in both shRNA RNA-Seq and eCLIP data, and removing eCLIP interactions from shRNA RNA-Seq ones, thus obtaining indirect interactions.

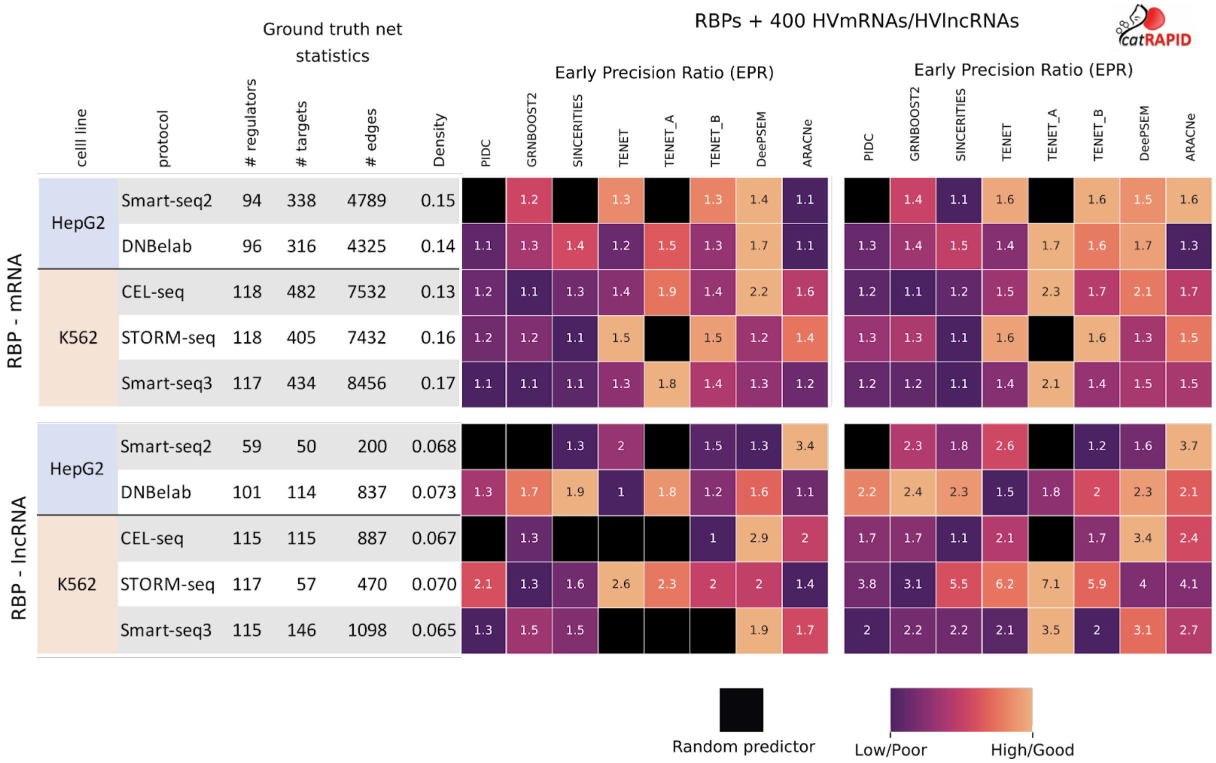


**Supplementary Figure S10**. **Related to Figure 3.** Heatmaps showing the EPR obtained for each scRNA-seq dataset (rows) and GRN inference method (columns) for the inference of RBP-mRNA (top) and RBP-lncRNA (bottom) interactions. A black box indicates EPR smaller than the one of a random predictor. The colors in the heatmap are scaled between 0 and 1 by row, ignoring values less than that of a random predictor. The table on the left of each heatmap shows the statistics of the cell-type specific eCLIP ground truth network for each dataset.

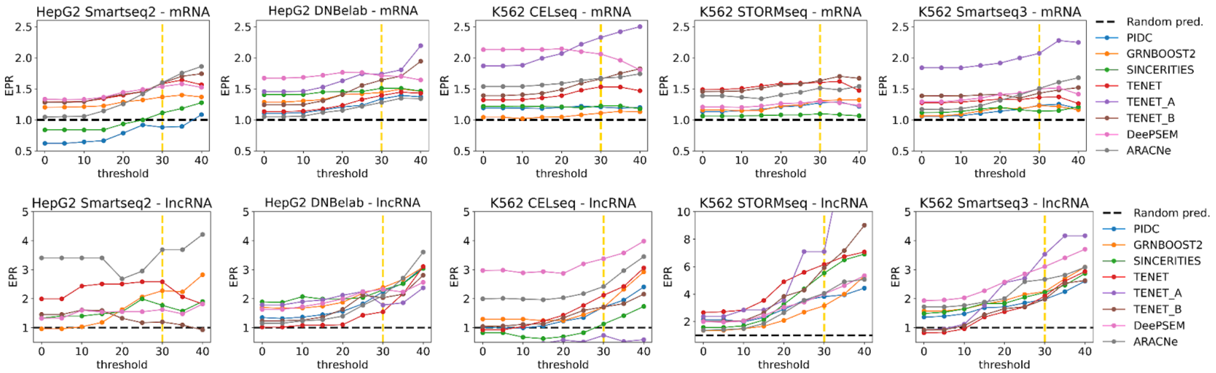


**Supplementary Figure S11**. **Related to Figure 3.** EPR as a function of the threshold on *cat*RAPID interaction propensity used for filtering the GRNs inferred by each algorithm. We show the datasets with 400 HVmRNAs in the top row and with 400 HVlncRNAs in the bottom row. The vertical yellow dashed line indicates the threshold selected for all the analyses. The horizontal black dashed line indicates the EPR of a random predictor. The y-axis range is different for the top and bottom rows, and in the bottom row the K562 STORM-seq dataset has a different range from the other datasets with HVlncRNAs. The ground truth network is given by eCLIP interactions.

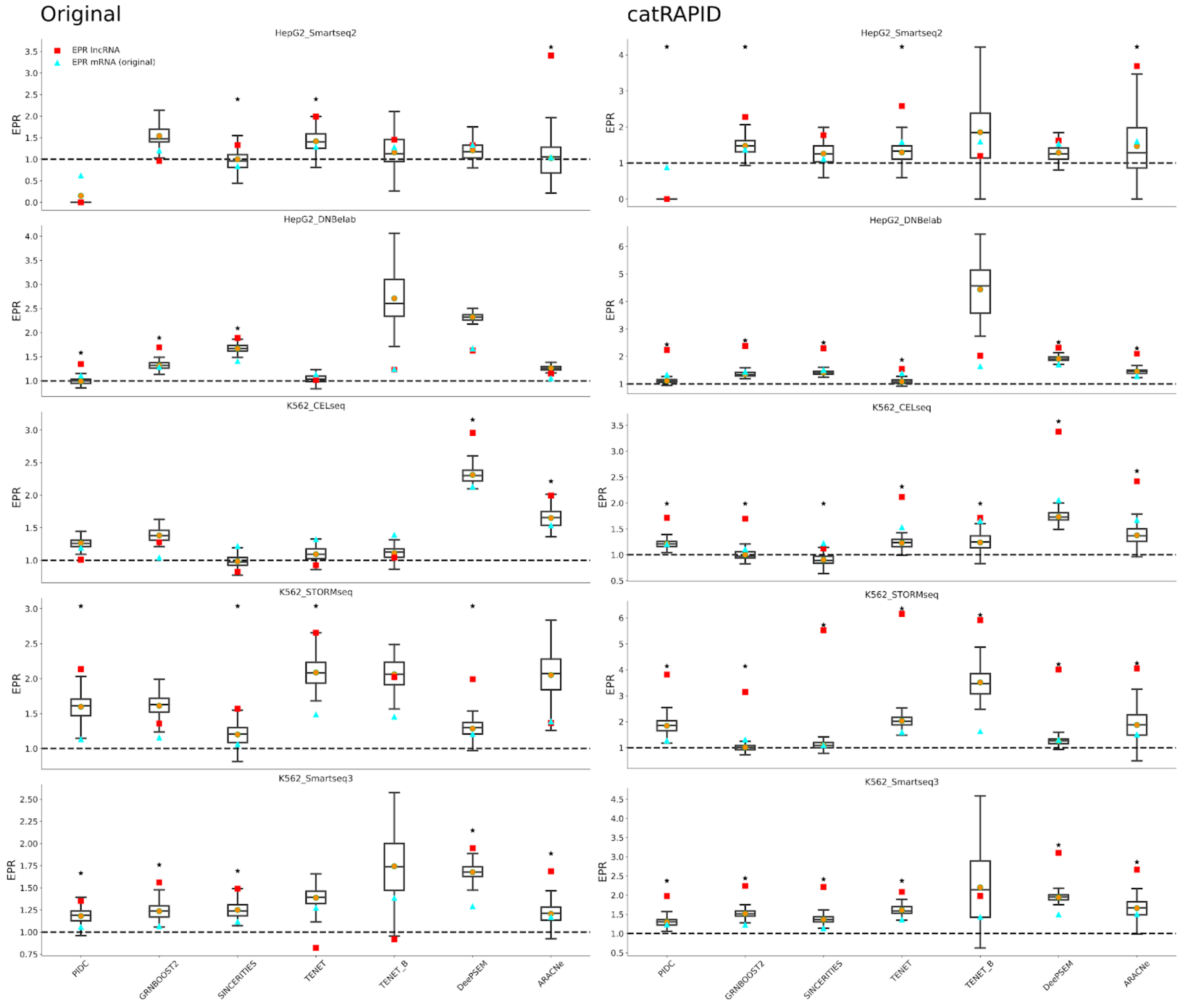


**Supplementary Figure S12**. **Related to Figure 3.** Downsampling of eCLIP RBP-mRNA networks for comparison of the inference performance between RBP-mRNA and RBP-lncRNA interactions. In each panel, the boxplots show the distribution of EPR values for the inferred RBP-mRNA interactions over 100 random samplings of the ground truth eCLIP RBP-mRNA network to match, on average, the number of RBPs, targets and edges of the corresponding RBP-lncRNA eCLIP network. The red square shows the EPR of the RBP-lncRNA interactions for the corresponding algorithm and dataset, the cyan triangle shows the EPR of the RBP-mRNA interactions before downsampling. The black star indicates a p-value smaller than 0.05, computed as the probability of sampling a value larger than the EPR of the RBP-lncRNA interactions from the Empirical Cumulative Distribution Function (ECDF) obtained from the 100 random samplings of the RBP-mRNA network (see **Supplementary Materials**). **Left**. Original rankings. **Right**. Rankings obtained after the *cat*RAPID-based filter.

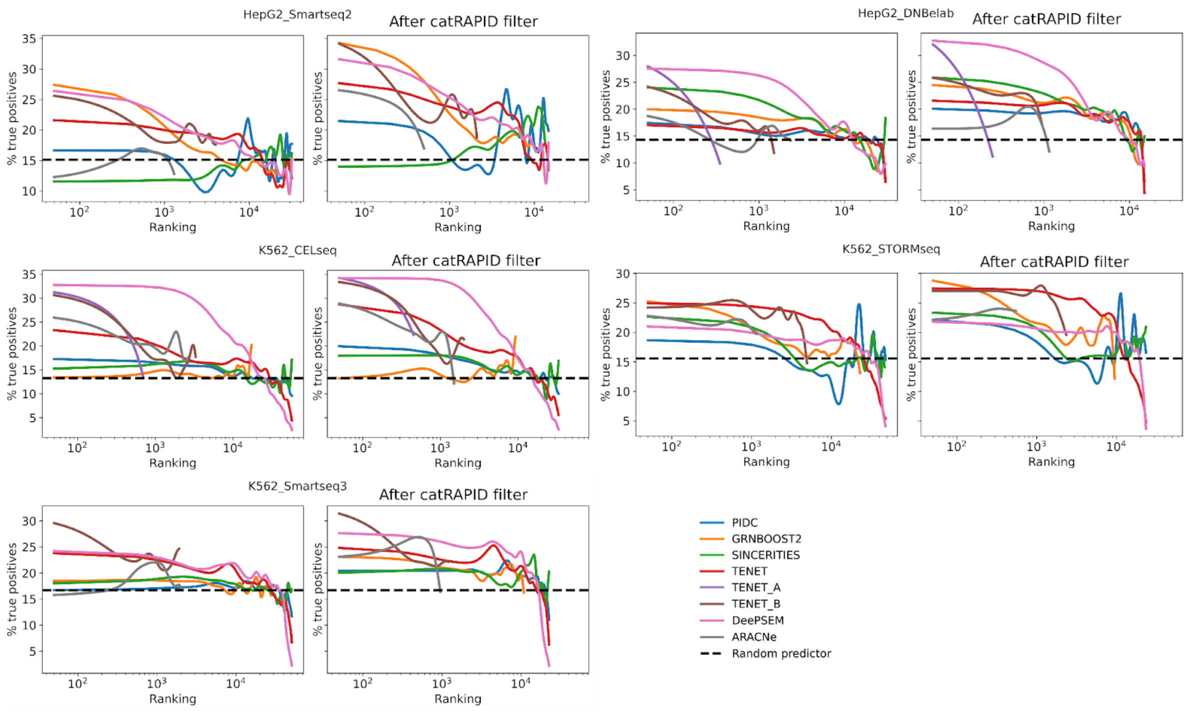


**Supplementary Figure S13**. **Related to Figure 3.** Percentage of true positive RBP-target interactions, according to the cell-type specific eCLIP ground truth network, as a function of the rank of the interactions inferred from scRNA-seq data, for each algorithm, before and after the *cat*RAPID-based filter, for the datasets with 400 HVmRNAs.

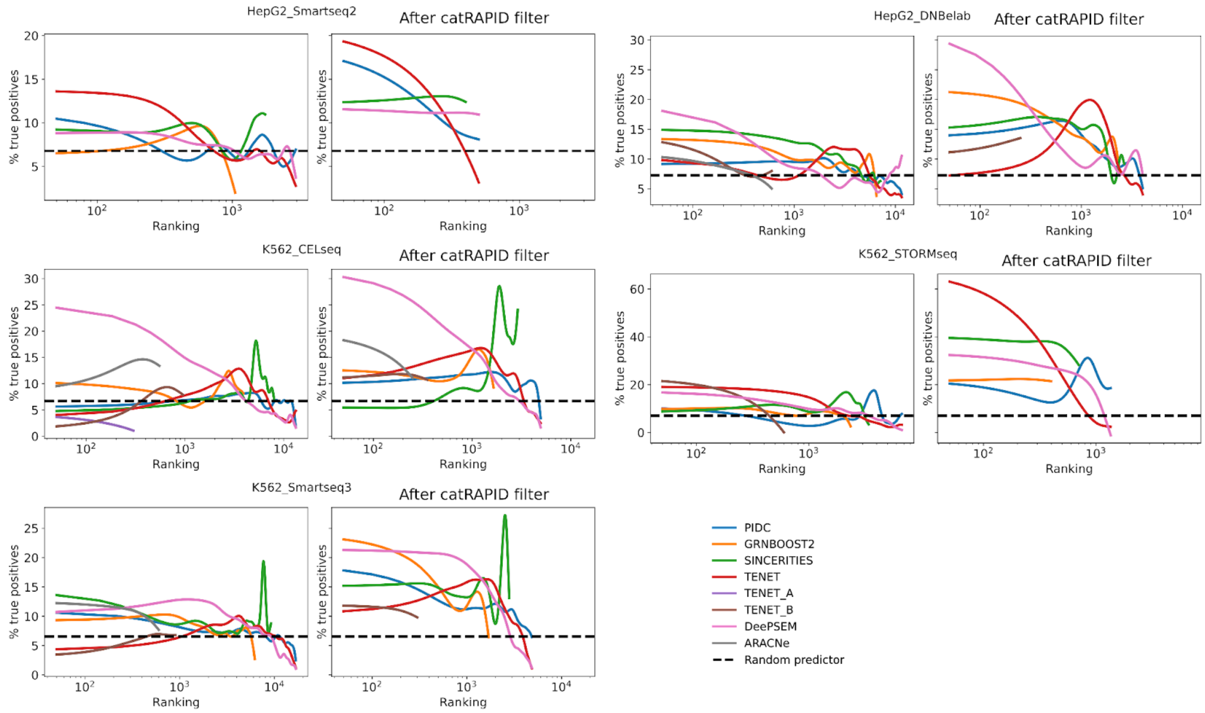


**Supplementary Figure S14**. **Related to Figure 3.** Percentage of true positive RBP-target interactions, according to the cell-type specific eCLIP ground truth network, as a function of the rank of the interactions inferred from scRNA-seq data, for each algorithm, before and after the *cat*RAPID-based filter, for the datasets with 400 HVlncRNAs.

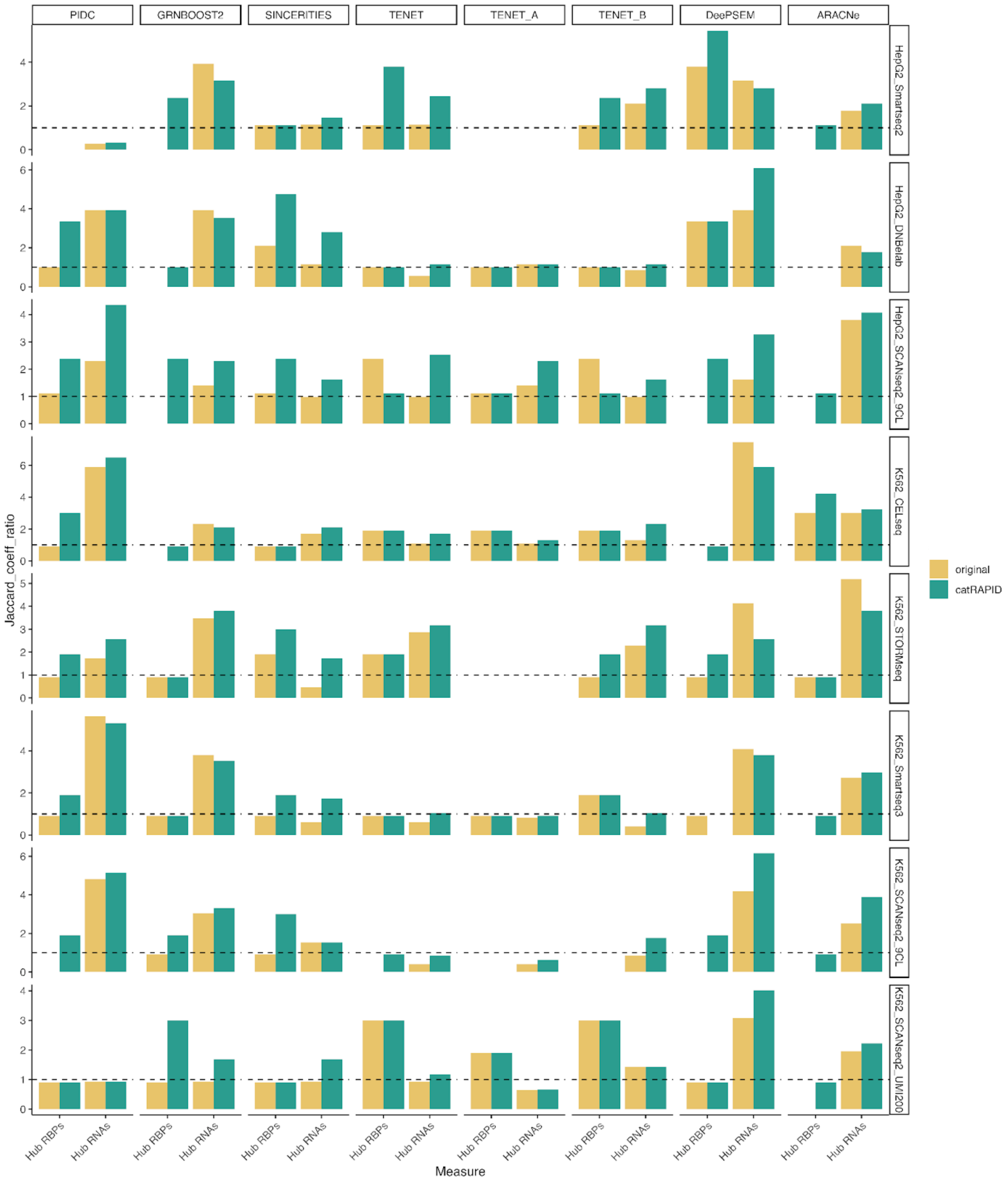


**Supplementary Figure S15. Related to Figure 4.** Bar plots showing the JCR for hub RBPs and hub RNAs before ("original") and after ("*cat*RAPID") the *cat*RAPID-based filter, for each scRNA-seq dataset (rows) with 500 HVGs and each GRN inference algorithm (columns). The dashed black line indicates the JCR of a random predictor.

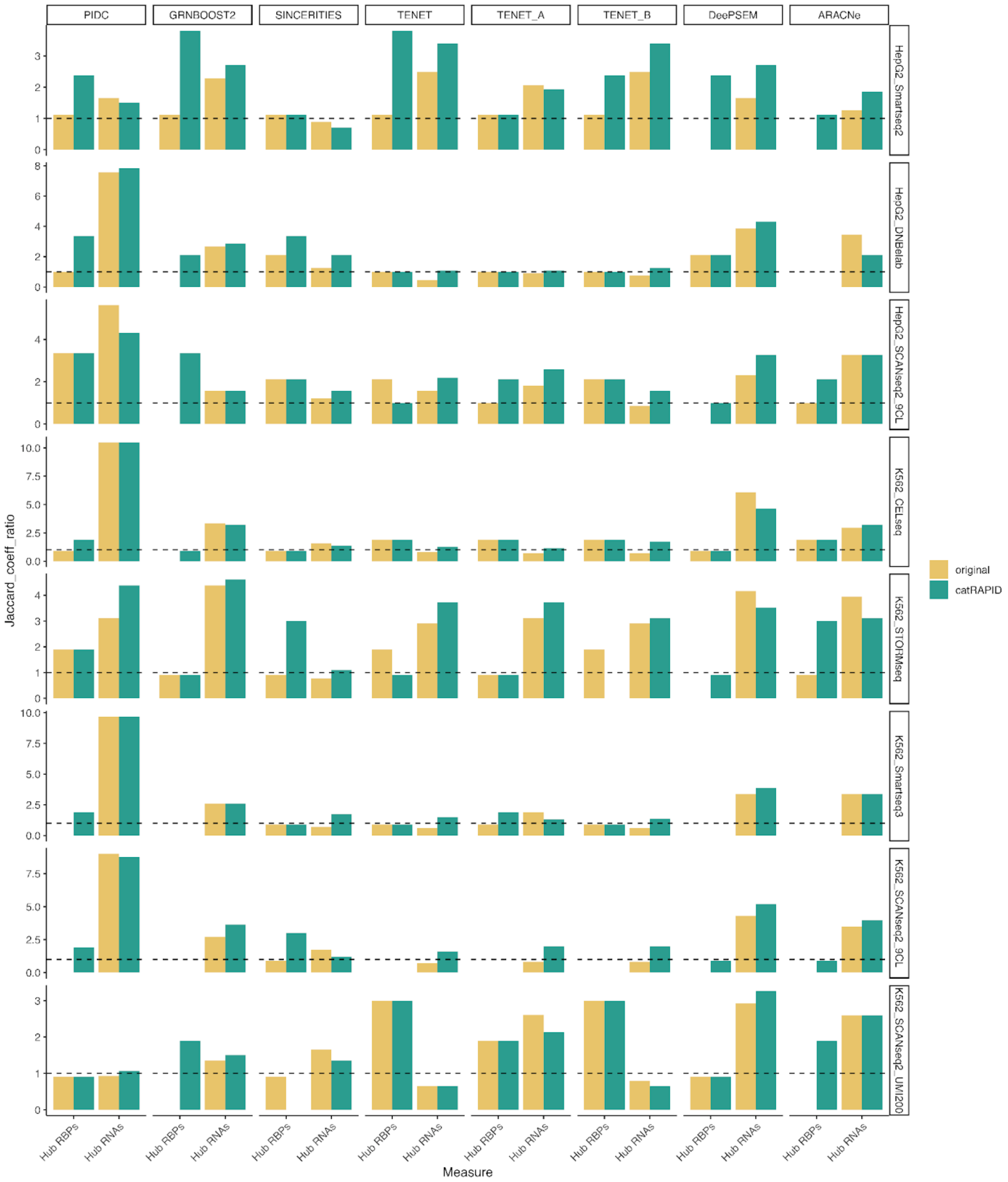


**Supplementary Figure S16. Related to Figure 4.** Bar plots showing the JCR for hub RBPs and hub RNAs before ("original") and after ("*cat*RAPID") the *cat*RAPID-based filter, for each scRNA-seq dataset (rows) with 1000 HVGs and each GRN inference algorithm (columns). The dashed black line indicates the JCR of a random predictor.

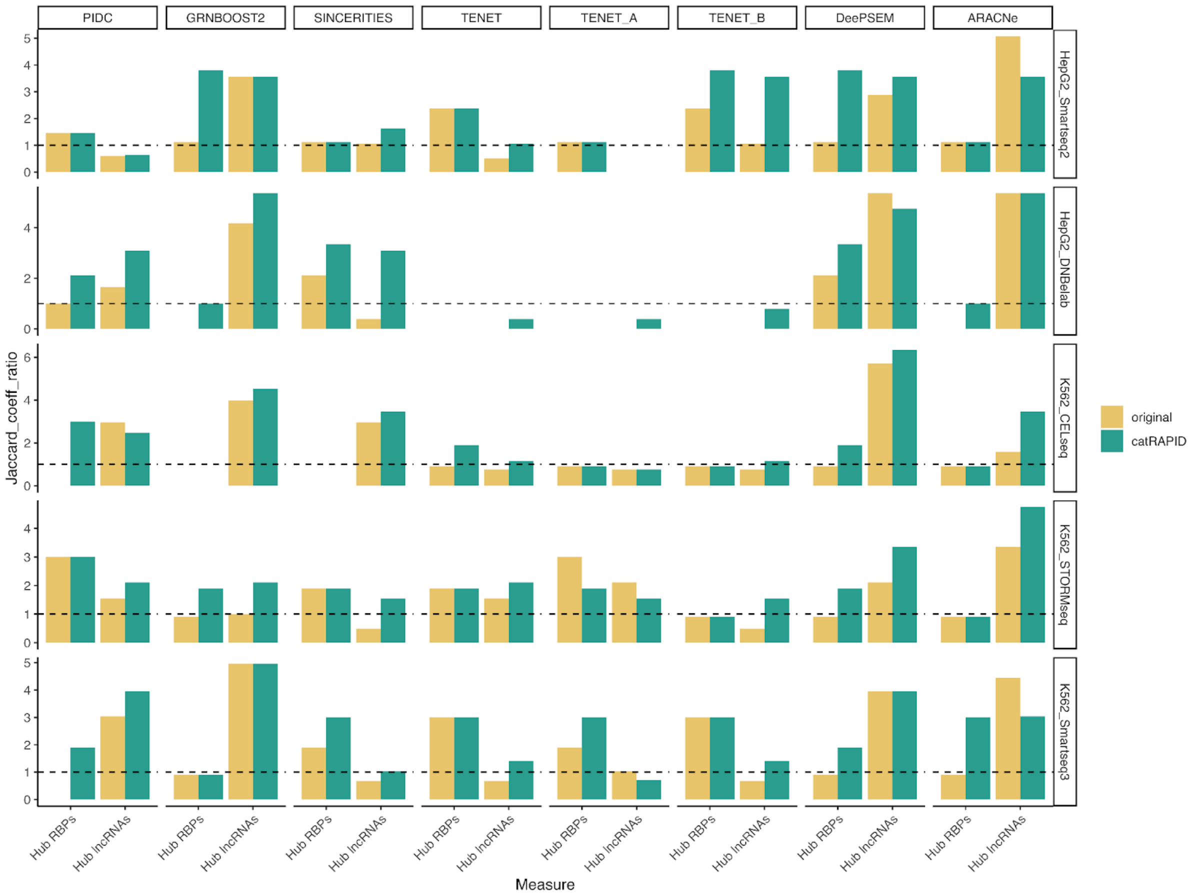


**Supplementary Figure S17. Related to Figure 4.** Bar plots showing the JCR for hub RBPs and hub lncRNAs before ("original") and after ("*cat*RAPID") the *cat*RAPID-based filter, for each scRNA-seq dataset (rows) with 400 HVlncRNAs and each GRN inference algorithm (columns). The dashed black line indicates the JCR of a random predictor.

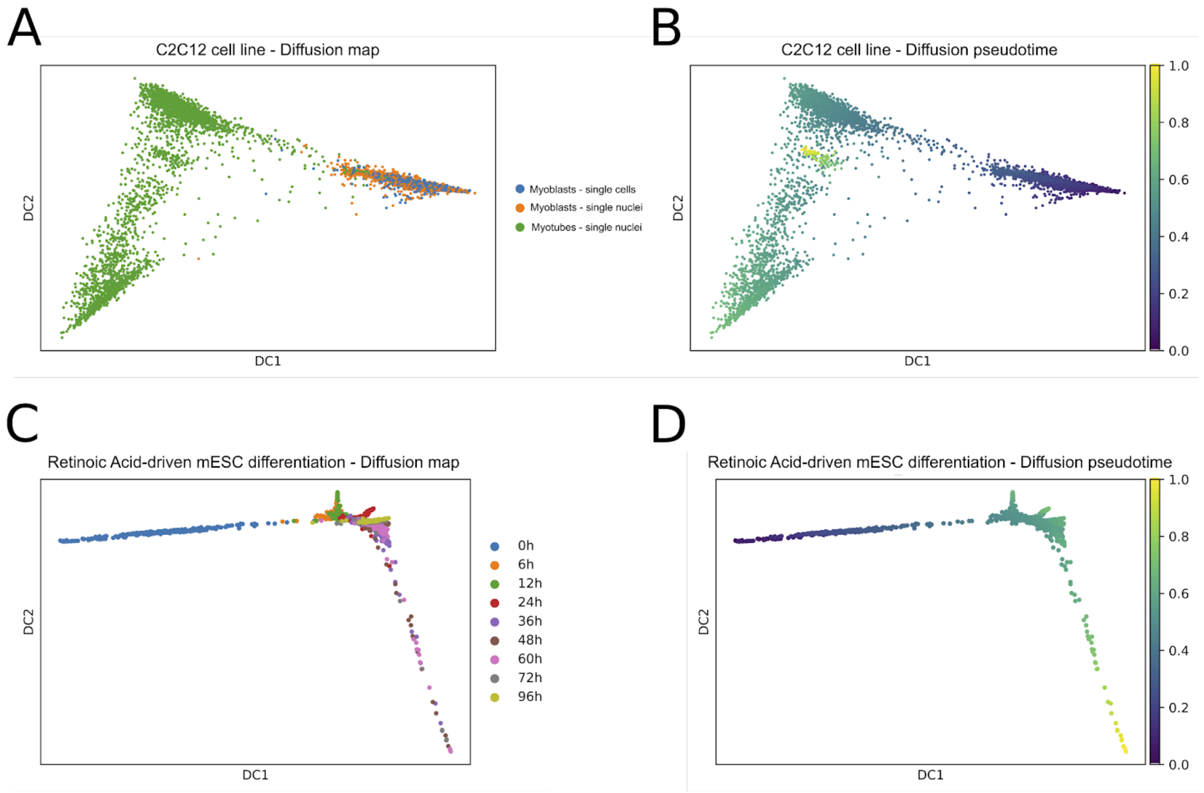


**Supplementary Figure S18. Related to Figure 5.** (**A-B**) Diffusion map and diffusion pseudotime calculated on the single-cell RNA-seq of myoblasts, the single-nuclei RNA-seq of myoblasts and the single-nuclei RNA-seq of myotubes from the SPLiT-seq dataset of the C2C12 murine cell line recapitulating myoblasts to myotubes differentiation. (**C-D**) Diffusion map and diffusion pseudotime calculated on the single-cell RNA-seq of mESCs sequenced at 9 time points, from 0 hours to 96 hours, of differentiation driven by Retinoic Acid.

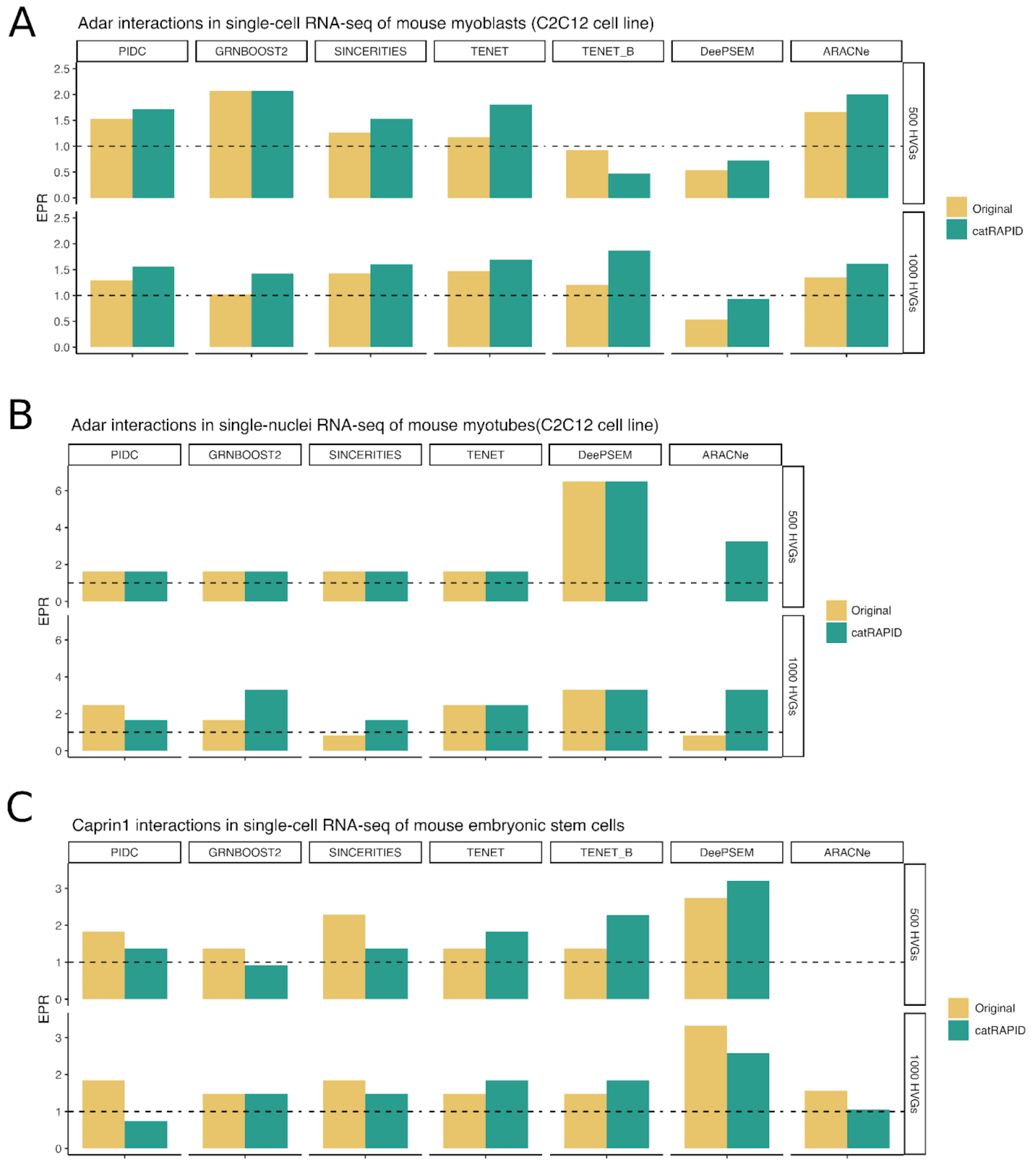


**Supplementary Figure S19. Related to Figure 5. A**) EPR measured for the inference made on the scRNA-seq of myoblasts from the SPLiT-seq dataset of the C2C12 murine cell line recapitulating myoblasts to myotubes differentiation. Performances are tested on the ADAR1 RIP-seq experiments at 0h. (**B**) EPR measured for the inference made on snRNA-seq of myotubes from the SPLiT-seq dataset of the C2C12 murine cell line recapitulating myoblasts to myotubes differentiation. Performances are tested on the ADAR1 RIP-seq experiments at 72h. (**C**) EPR measured for the inference made on the scRNA-seq of mouse embryonic stem cells (mESCs) from the SCRB-seq dataset. Performances are tested on the Caprin1 RIP-seq experiments at 0h. In all panels we show only algorithms for which at least two experimental interactions are present in the inferred network, before the *cat*RAPID-based filter.

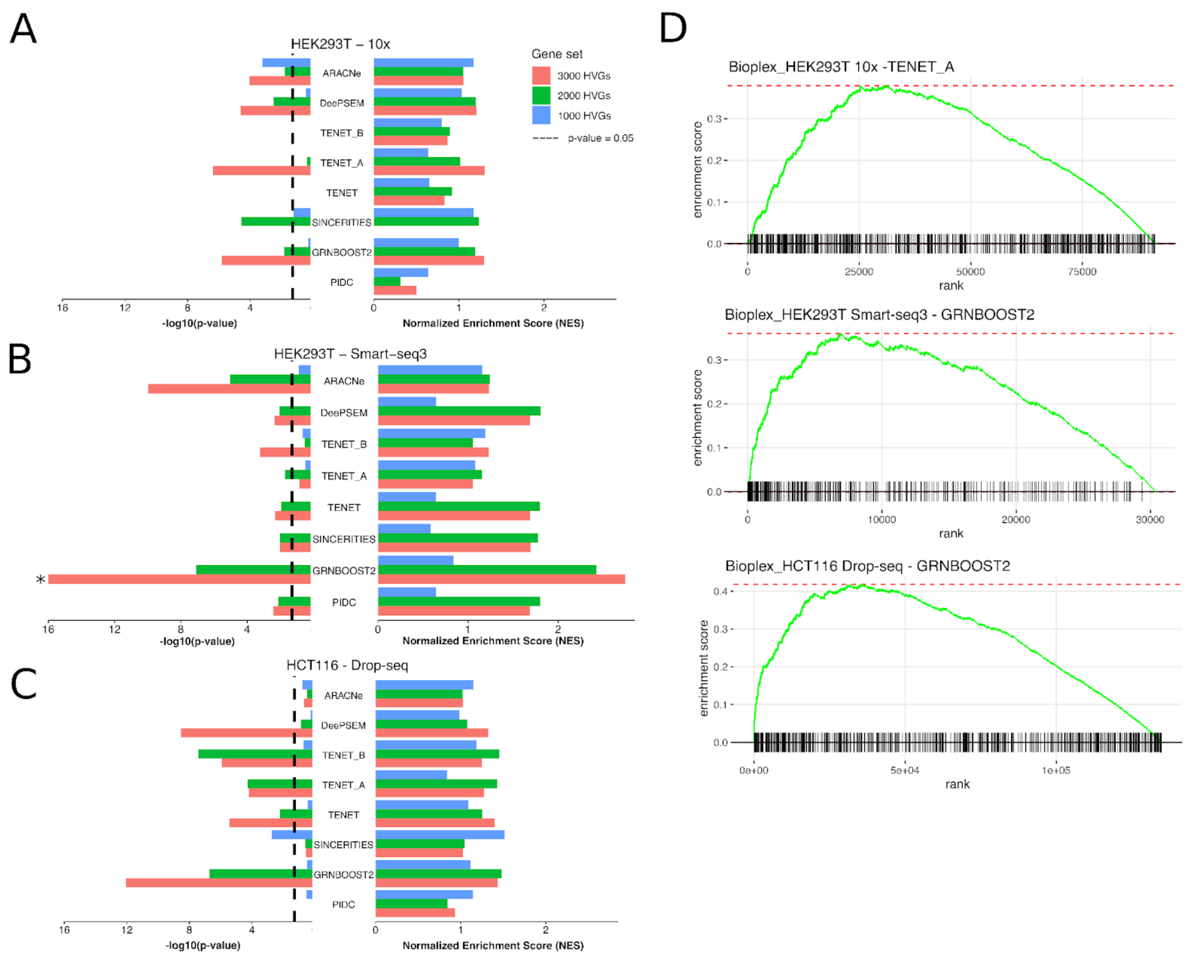


**Supplementary Figure S20. Related to Figure 6.** (**A-B-C).** Bar plots showing the -log10(p-value) (left) and Normalized Enrichment Score (NES) obtained from the Gene Set Enrichment Analysis (GSEA) on the inferred RBP-RBP pairs ranked according to the fraction of shared RNA targets in *cat*RAPID-based filtered rankings, for each GRN inference method. (**A**) scRNA-seq of the HEK293T cell line obtained with the 10x protocol. (**B**) scRNA-seq of the HEK293T cell line obtained with the Smart-seq3 protocol. (**C**) scRNA-seq of the HCT116 cell line obtained with the Drop-seq protocol. p-values capped to $10^{-16}$ are indicated by a star. (**D**) GSEA enrichment plots for the most significant inference method for each scRNA-seq dataset (3000 HVGs).
